# Characterizing smoking-induced transcriptional heterogeneity in the human bronchial epithelium at single-cell resolution

**DOI:** 10.1101/484394

**Authors:** Grant E. Duclos, Vitor H. Teixeira, Patrick Autissier, Yaron B. Gesthalter, Marjan A. Reinders-Luinge, Robert Terrano, Yves M. Dumas, Gang Liu, Sarah A. Mazzilli, Corry-Anke Brandsma, Maarten van den Berge, Sam M. Janes, Wim Timens, Marc E. Lenburg, Avrum Spira, Joshua D. Campbell, Jennifer Beane

## Abstract

The human bronchial epithelium is composed of multiple, distinct cell types that cooperate to perform functions, such as mucociliary clearance, that defend against environmental insults. While studies have shown that smoking alters bronchial epithelial function and morphology, the precise effects of this exposure on specific cell types are not well-understood. We used single-cell RNA sequencing to profile bronchial epithelial cells from six never- and six current smokers. Unsupervised analyses identified thirteen cell clusters defined by unique combinations of nineteen distinct gene sets. Expression of a set of toxin metabolism genes localized to ciliated cells from smokers. Smoking-induced airway remodeling was characterized by a loss of club cells and extensive goblet cell hyperplasia. Finally, we identified a novel peri-goblet epithelial subpopulation in smokers that expressed a marker of bronchial premalignant lesions. Our data demonstrates that smoke exposure drives a complex landscape of cellular and molecular alterations in the human bronchial epithelium that may contribute to the onset of smoking-associated lung diseases.

---

The human bronchus is lined with a pseudostratified epithelium that acts as a physical barrier against exposure to harmful environmental insults such as inhaled toxins, allergens, and pathogens^1-2^. The bronchial epithelium is a complex tissue, predominantly composed of ciliated, goblet, club, and basal epithelial cells. These cell types cooperate to perform mucociliary clearance, which is the process that mediates the capture and removal of inhaled substances^1-2^. Goblet cells secrete components of a mucosal lining that entraps inhaled particulate matter, which is propelled out of the airways by mechanical beating of ciliated cells^1-2^. Club cells also perform a secretory function^3^ and serve as facultative progenitors^4^, whereas basal cells are multipotent progenitors responsible for normal tissue homeostasis^5-7^. Interplay amongst these cells is required for proper function and long-term maintenance of the bronchial epithelium, but exposure to substances, like tobacco smoke, might alter or injure specific cell types and lead to tissue-wide dysfunction.

Inhalation of tobacco smoke exposes the bronchial epithelium to toxins, carcinogens, and free radicals^8-11^, but cellular injuries and abnormalities associated with this exposure are complex and have not been fully characterized. Previous studies have described smoking-induced epithelial changes, such as increased goblet cell numbers^12-14^ and reduced ciliary length^15-16^. Robust transcriptomic alterations have also been observed in the bronchial epithelium of smokers, involving the up-regulation of genes linked to xenobiotic metabolism and the oxidative stress response^17-18^. Furthermore, it has been reported that a subset of gene expression alterations detected in smokers persists years after smoking cessation^18^. However, the aforementioned transcriptomic studies profiled bronchial tissue in “bulk”, masking cell type-specific contributions to the smoking-associated gene expression signature.

To overcome the limitations of bulk tissue analyses, we used single-cell RNA sequencing (scRNA-Seq) to profile the transcriptomes of individual bronchial cells from healthy never and current smokers. We identified bronchial subpopulations using an unsupervised machine learning algorithm then immunostained bronchial tissue from independent cohorts of never and current smokers to validate robust smoking-associated findings. In the airways of smokers, we described a metabolic response specific to ciliated cells, a shift in the presence of club and goblet cells, and the emergence of a previously uncharacterized epithelial subpopulation.

## RESULTS

### Single-cell RNA-Seq was used to identify bronchial subpopulations in the airways of never and current smokers

Bronchial brushings were procured by bronchoscopy from the right mainstem bronchus of 6 healthy current smokers and 6 healthy never smokers (**Supplementary Table 1**) and single ALCAM^+^ epithelial cells^19^ and CD45^+^ white blood cells (WBCs) were isolated from each donor (Figure 1a, **Supplementary Figure 1)**. The CEL-Seq single-cell RNA sequencing (scRNA-Seq) protocol^20^ was used to profile the transcriptomes of 84 epithelial cells and 11 WBCs from each of the 12 donors (1,140 total cells: 1,008 epithelial cells and 132 WBCs). Low quality cells were excluded from downstream analyses, leaving 796 cells (753 epithelial cells, 43 WBCs) (**Supplementary Figure 2-3**) expressing an average of 1,817 genes per cell. Expression of known marker genes for bronchial cell types were detected in largely non-overlapping cells, including KRT5 for basal cells, FOXJ1 for ciliated cells, SCGB1A1 for club cells, MUC5AC for goblet cells, and CD45 for white blood cells (Figure 1b). Given the relatively small number of subjects, we sought to determine if smoking-associated gene expression changes identified in these donors reflected those observed in a larger, independent cohort of never- and current smokers. Data from all cells procured from each donor was combined to generate *in silico* “bulk” bronchial brushings. Analysis of differential expression between never- and current smoker *in silico* bulk samples revealed associations that were highly correlated (Spearman’s *r* = 0.45) with those observed in a previously published bulk bronchial brushing dataset generated by microarray^18^ (**Supplementary Figure 4**).

**Figure 1.**
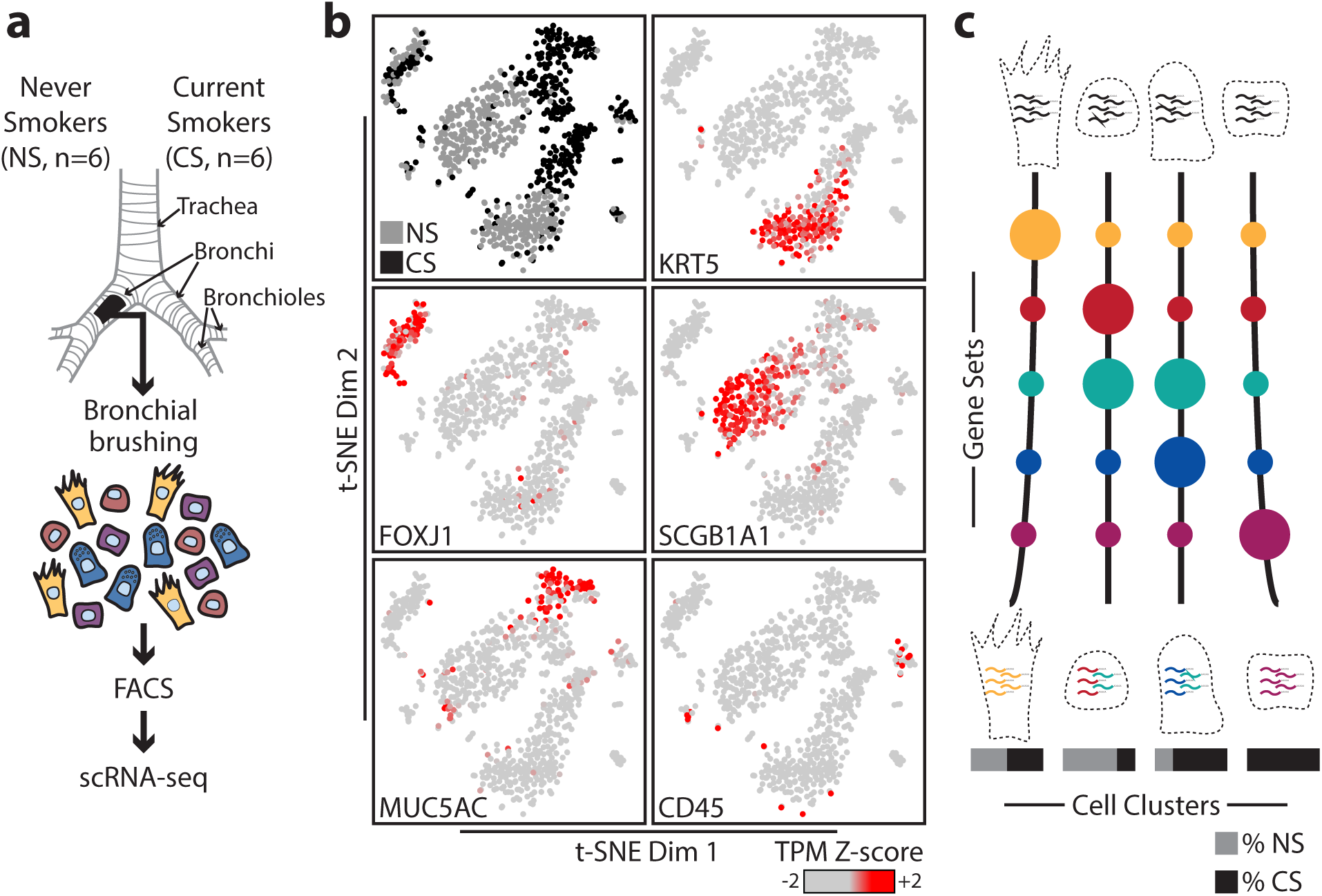
Single-cell RNA sequencing of human bronchial cells from never and current smokers. (**a**) Bronchial brushings were procured from the right mainstem bronchus of 6 never smokers and 6 current smokers. Bronchial tissue was dissociated, single cells were isolated by FACS, and single-cell RNA libraries were prepared and sequenced. (**b**) t-Distributed Stochastic Neighbor Embedding (t-SNE) was performed to illustrate transcriptomic relationships amongst cells. Donor smoking status (NS = never smoker, CS = current smoker) was visualized for each cell as well as expression of bronchial cell type marker genes (z-normalized transcripts per million (TPM) values) across all cells: KRT5 (basal), FOXJ1 (ciliated), SCGB1A1 (club), MUC5AC (goblet), and CD45 (WBC). (**c**) An unsupervised analytical approach (LDA) was utilized to identify distinct cell clusters and sets of co-expressed genes. Cell clusters were defined by unique gene set expression patterns and never- or current smoker cell enrichment was assessed.

In order to characterize cellular subpopulations beyond known cell type markers, we utilized Latent Dirichlet Allocation (LDA) as an unsupervised framework to assign cells to clusters and to identify distinct sets of co-expressed genes across all cells (Figure 1C). LDA divided the dataset into 13 distinct cell clusters and 19 sets of co-expressed genes (Figure 2a-b, **Supplementary Figure 5-8**). Each cell cluster was defined by the expression of a unique combination of gene sets and each gene set was defined by a unique expression pattern amongst clusters (Figure 2a-b, **Supplementary Figure 9**). Cell types were defined for 8 of the 13 clusters based on medium to high marker gene expression: cell clusters C-2 and C-4 expressed KRT5, C-5 and C-11 expressed FOXJ1, C-1 and C-8 expressed SCGB1A1, and C-3 expressed MUC5AC (Figure 2c). Cluster C-7 expressed WBC marker CD45 (Figure 2c) and Fisher’s exact test (FET) was used to show that C-7 was enriched with sorted CD45^+^ cells (FET *p* = 9.6 x 10^−47^). C-7 cells also expressed several T cell receptor genes (e.g. TRBC2, TRGC1), indicating a T cell lineage (**Supplementary Figure 10**). Low levels of SCGB1A1 transcripts were detected in cluster C-10 (SCGB1A1^low^) and CFTR was expressed by cluster C-13, which suggests that these cells may be ionocytes^21^ (**Supplementary Figure 11**). Marker gene expression was not detected in clusters C-6, C-9, and C-12 (Figure 2c). Enrichment of current smoker cells was observed in goblet cell cluster C-3, as well as C-9 and C-12, whereas enrichment of never smoker cells was observed in club cell cluster C-1, SCGB1A1^low^ cluster C-10 and basal clusters C-2 and C-4 (Figure 2d). Furthermore, a subset of gene sets expressed by specific clusters of ciliated-, club-, goblet-, and basal cells, as well as those without a cell type designation, were differentially expressed between never- and current smokers in transcriptomic data generated from bulk bronchial tissue^18^ (Figure 2a-b, **Supplementary Figure 12**). Therefore, smoking-induced gene expression changes reported in bulk tissue are likely driven by alterations to multiple bronchial cell types.

**Figure 2.**
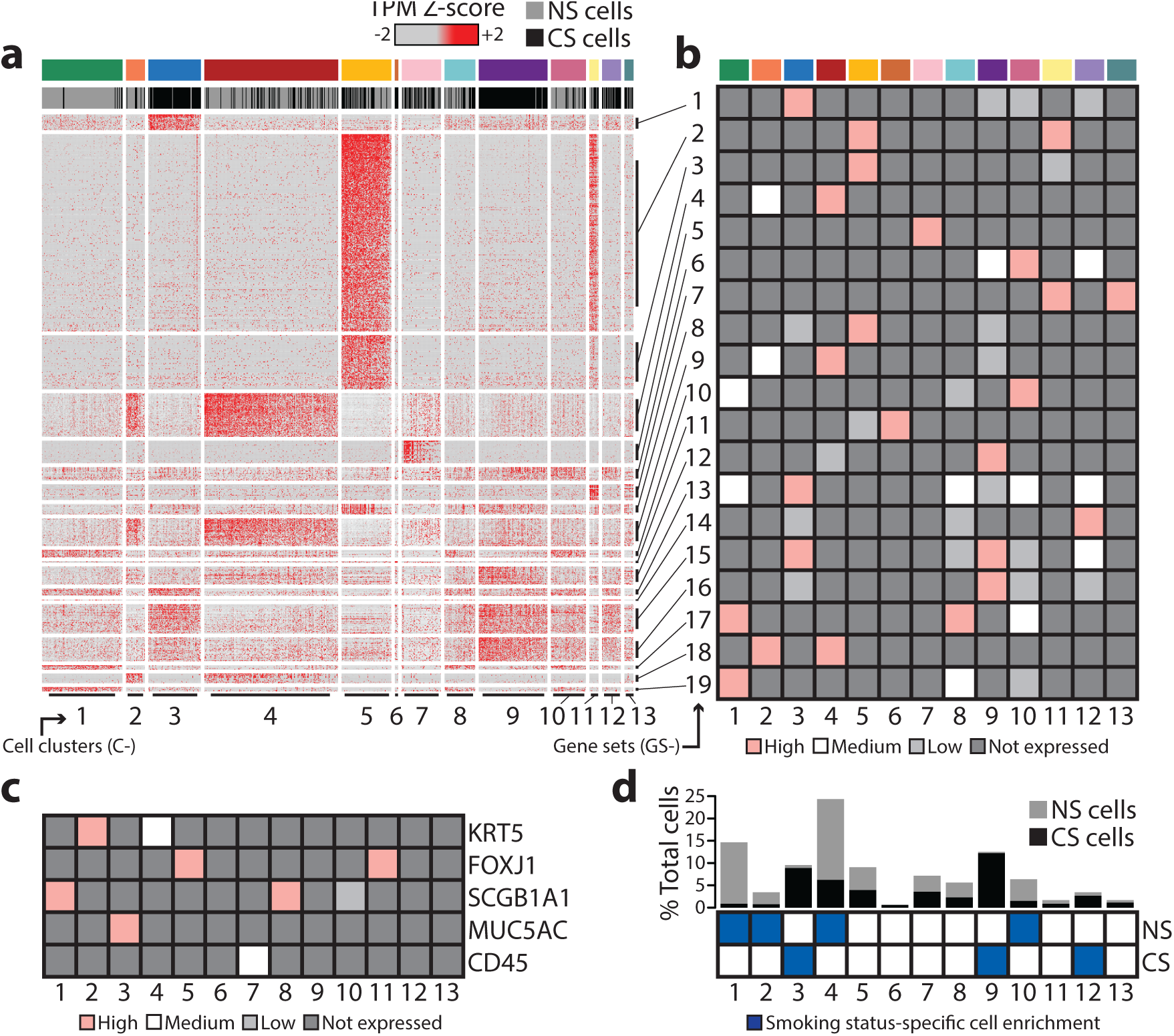
Characterization of bronchial cluster transcriptomic profile, cell type, and smoking status. (**a**) Global transcriptomic profiles of 13 bronchial cell clusters were defined by expression of unique combinations of 19 gene sets and visualized by heatmap (z-normalized TPM values). (**b**) A MetaGene was generated for each gene set (GS-1 to 19) and mean cluster-specific expression was designated: high (pink), medium (white), low (light grey), or not expressed (dark grey). (**c**) Mean expression of marker genes was summarized for each cluster designated: high (pink), medium (white), low (light grey), or not expressed (dark grey). (**d**) Per cluster percentage of total cells and the ratio of never-(NS) and current (CS) smoker cells was calculated. Fisher’s exact test (FET) was used to assess per cluster statistical enrichment of NS or CS cells.

### Ciliated cell subpopulations and smoking-induced detoxification

We characterized transcriptomic similarities and differences amongst FOXJ1^+^ clusters C-5 and C-11 in order to define ciliated cell subpopulations detected in never and current smokers. Our data revealed that both clusters of ciliated cells expressed gene set GS-2, but could be differentiated based on expression of gene set GS-3 by cluster C-5 and gene set GS-7 by cluster C-11 (Figure 3a). GS-2 contains FOXJ1, in addition to genes involved with ciliary assembly, maintenance, and function, such as motor protein genes (e.g. DYNLL1, DNAH9) and intraflagellar transport genes (e.g. IFT57, IFT172) (Figure 3a, **Extended Supplementary Table 3**). GS-2 also includes anti-oxidant genes (e.g. PRDX5, GPX4, GSTA2), known transcriptional regulators of ciliogenesis (e.g. RFX2^22-23^, RFX3^24-25^), and surface proteins not previously attributed to ciliated cells (e.g. CDHR3, CD59). GS-3 also contains genes with known roles in airway ciliary biology, such as IFT88^26-28^ (required for ciliary formation) and DNAH5^29-31^ (required for ciliary motility). By contrast, gene set GS-7 is enriched with cell cycle-associated genes (**Extended Supplementary Table 3**), such as CDK1 and CCNB1 (G1/S transition) and TOP2A (S-phase DNA replication), as well as the transcription factor, HES6. Therefore, clusters C-5 and C-11 likely represent functionally distinct subpopulations of FOXJ1^+^ ciliated cells.

**Figure 3.**
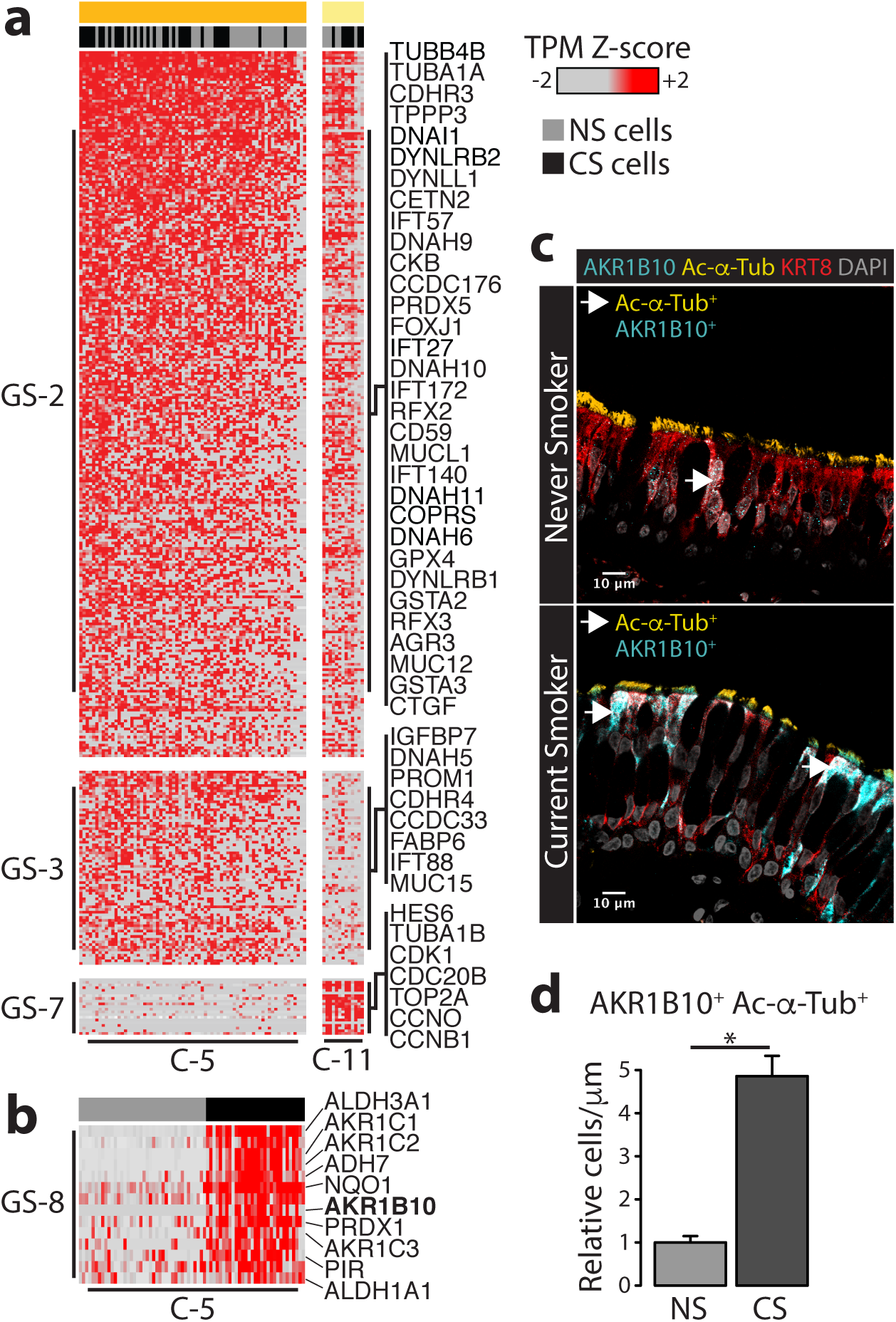
A smoking-induced detoxification program was observed in ciliated cells. (**a**) Expression of gene sets GS-2, GS-3, and GS-7 in clusters C-5 and C-11 was visualized by heatmap (z-normalized TPM values). (**b**) Cluster C-5 was split into never and current smoker subsets and expression of GS-8 genes was visualized by heatmap. (**c**) Bronchial tissue procured from an independent cohort of never and current smokers (UMCG Cohort, **Supplementary Table 2**) was immunostained for AKR1B10, Acetylated alpha tubulin (Ac-α-Tub), and KRT8. Representative images of never smoker (left) and current smoker (right) tissue were displayed. Arrows indicate ciliated cell (Ac-α-Tub^+^)- specific expression of AKR1B10. (**d**) Changes in tissue length (μm)-normalized numbers of AKR1B10^+^ Ac-α-Tub^+^ cells in current smokers, relative to never smokers, were assessed by WRS test.

We found that ciliated cells from current smokers expressed a distinct transcriptional signature. Specifically, the current smoker subset of cluster C-5 FOXJ1^+^ cells expressed gene set GS-8, which was enriched with genes encoding enzymes implicated in aldehyde and ketone metabolism, such as ALDH3A1, AKR1C1, and AKR1B10 (Figure 3b). This finding suggested that the gene expression response to toxic aldehydes and ketones present in tobacco smoke^8-9^ might be restricted to ciliated epithelial cells. To confirm that this set of enzymes localized to ciliated cells, we immunostained bronchial tissue procured from an independent cohort of never and current smokers (UMCG Cohort, **Supplementary Table 2**) for the aldo-keto reductase AKR1B10, as well as cilia-specific acetylated alpha tubulin (Ac-α-Tub) and the luminal cytokeratin, KRT8, which is expressed by all non-basal cells (Figure 3c). We found that AKR1B10 was robustly expressed in the airways of current smokers and numbers of AKR1B10^+^ ciliated cells were significantly higher than those observed in never smokers (*p* = 7.4 x 10^−7^; Figure 3c-d). AKR1B10 was detected throughout the cytoplasm of smoker ciliated cells, as well as at the base of the cilia (Figure 3c). AKR1B10^+^ ciliated cells were uncommon in never smokers and overall low magnitude of AKR1B10 expression was observed in these cells (Figure 3c). We detected rare instances of non-ciliated AKR1B10^+^ KRT8^+^ cells (**Supplementary Figure 13a**), but AKR1B10^+^ KRT8^−^ cells were not observed. We also confirmed that AKR1B10 was not expressed by current smoker MUC5AC^+^ goblet cells (**Supplementary Figure 13b**). Overall, these results demonstrate that ciliated cells express a specific set of detoxification genes in response to smoke exposure.

### Club cell depletion and goblet cell expansion in the airways of smokers

Our data revealed that the largest cluster of SCGB1A1^+^ cells, C-1, was enriched with never smoker cells (Figure 2d), indicating that this subpopulation of club cells was depleted from the airways of smokers. C-1 cells distinctly expressed high levels of gene set GS-19, which contains MUC5B, in addition to SCGB3A1 and transcription factors TCF7, FOS, and JUN (Figure 4a). However, SCGB1A1 (included in gene set GS-17) was also highly expressed by cluster C-8, which was not affected by smoking status (Figure 2d). Therefore, these results indicate that smoking is associated with a decrease in MUC5B^+^ SCGB1A1^+^ (C-1) club cell content. Furthermore, gene set GS-13, which contains immunologically relevant genes BPIFB1^32^ and PIGR^33^ (Figure 4a), was expressed by SCGB1A1^+^ cells (C-1 and C-8) as well as MUC5AC^+^ cluster C-3, indicating that there may be functional overlap amongst club and goblet cells.

**Figure 4.**
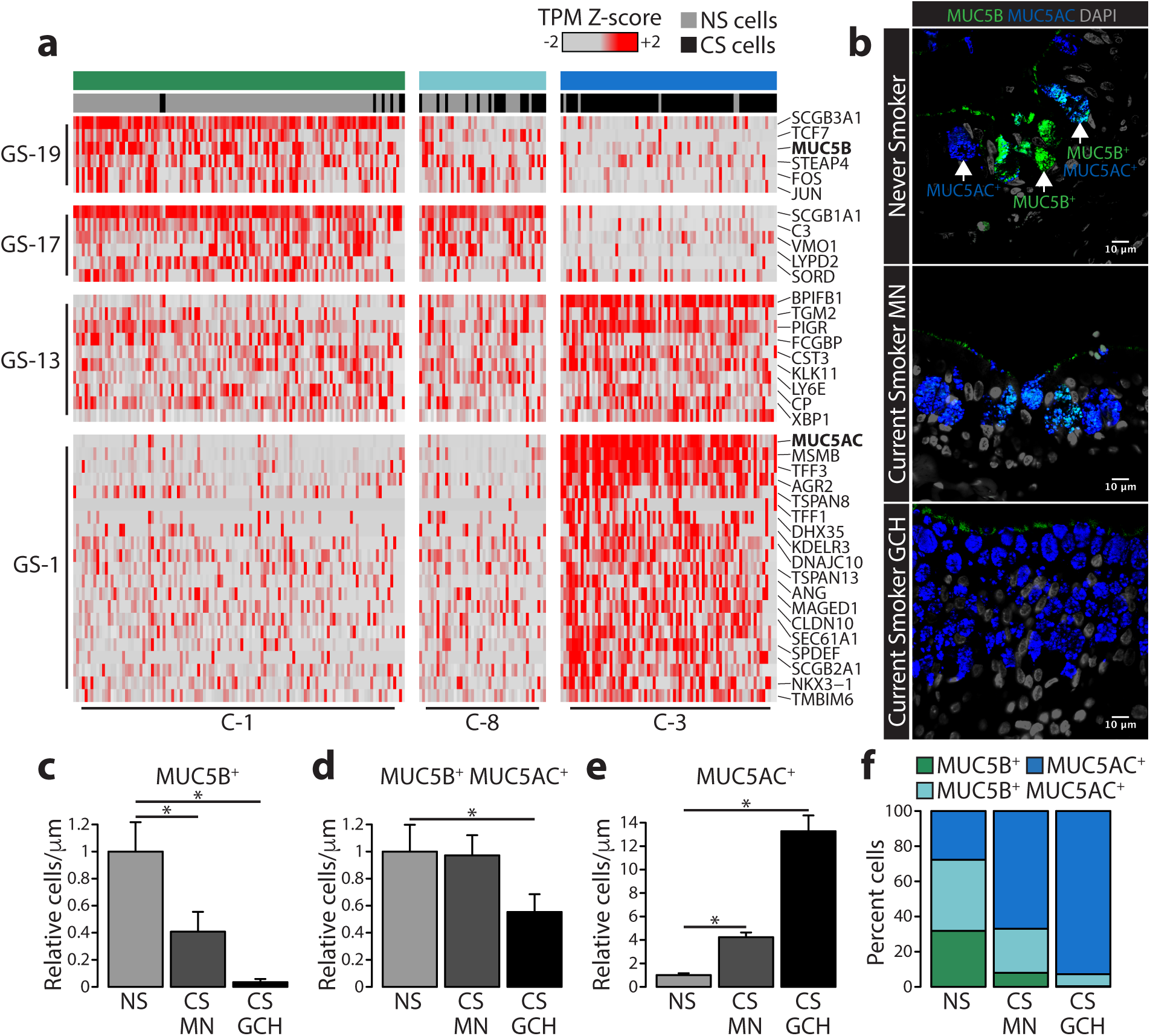
Smoking is associated with increased numbers of goblet cells and decreased numbers of club cells in the bronchial epithelium. (**a**) Expression of gene sets GS-19, GS-17, GS-13, and GS-1 in clusters C-1, C-8, and C-3 was visualized by heatmap (z-normalized TPM values). Bronchial tissue procured from an independent cohort of never and current smokers (UMCG Cohort, **Supplementary Table 2**) was immunostained for MUC5B and MUC5AC. (**b**) Representative images of never smoker tissue (left), morphologically normal (MN) current smoker tissue (middle), and current smoker goblet cell hyperplasia (GCH) (right) were displayed. Changes in tissue length (μm)-normalized numbers of (**c**) MUC5B^+^ (**d**) MUC5B^+^ MUC5AC^+^, and (**e**) MUC5AC^+^ cells in current smoker morphologically normal and GCH tissue were assessed relative to never smokers by WRS test. (**f**) Proportions of MUC5B^+^, MUC5B^+^ MUC5AC^+^, and MUC5AC^+^ cells were examined in never smokers, as well as MN and GCH current smoker tissue.

The MUC5AC^+^ goblet cell cluster C-3 was significantly enriched with current smoker cells (Figure 2d), which is consistent with prior studies showing that smoking is associated with increased bronchial goblet cell abundance^12-14^. Cluster C-3 expressed gene set GS-1, which contains the goblet cell marker gene, MUC5AC, as well as several genes with known roles in goblet cell biology, such as SPDEF^34^, AGR2^35^, and TFF3^36^ (Figure 4a). Genes associated with the unfolded protein response (UPR) are present in GS-1 (e.g. KDLER3, DNAJC10) (**Extended Supplementary Table 3**). We also identified several unique goblet cell surface markers (e.g. CLDN10, TSPAN8, TSPAN13), as well as a transcription factor (NKX3-1) whose role in the goblet cell transcriptional program is unknown (Figure 4a). Therefore, this data indicates that smoking is associated with increased numbers of MUC5AC^+^ goblet cells.

To confirm smoking-associated shifts in club and goblet cell numbers, bronchial tissue procured from an independent cohort of never and current smokers (UMCG Cohort, **Supplementary Table 2**) was immunostained for markers of club (MUC5B) and goblet (MUC5AC) cells (Figure 4b). Imaging data revealed cell subpopulations that exclusively express MUC5B or MUC5AC, as well as those that co-express both MUC5B and MUC5AC (Figure 4b). The airways of never smokers contained similar numbers of MUC5B^+^, MUC5B^+^ MUC5AC^+^, and MUC5AC^+^ cells (Figure 4b,f). The bronchial epithelium of current smokers, however, took on two distinct phenotypes: tissue regions described as “morphologically normal” (MN), which were similar to never smokers, and regions characterized by high MUC5AC^+^ cell density, referred to as goblet cell hyperplasia (GCH) (Figure 4b). In the MN smoker tissue, we observed a significant decrease in MUC5B^+^ cells (*p* = 0.02) (Figure 4c) and a significant increase in MUC5AC^+^ cells (*p* = 1.5 x 10^−6^) (Figure 4e), relative to never smokers, but no change in MUC5B^+^ MUC5AC^+^ content was observed (Figure 4d). Differences between smoker GCH and never smoker epithelium, however, were more pronounced. Near complete loss of MUC5B^+^ cells was observed in smoker GCH (*p* = 1.8 x 10^−5^; Figure 4c), along with a significant loss of MUC5B^+^ MUC5AC^+^ cells (*p* = 0.02; Figure 4d), relative to never smokers. GCH-associated alterations were accompanied by a 13-fold increase in MUC5AC^+^ cells (*p* = 7.4 x 10^−7^; Figure 4e-f). Additional immunostaining for KRT5 expression in the same bronchial tissue revealed that basal cell content was not affected by smoking status and did not vary between MN and GCH regions (**Supplementary Figure 14**). Overall, these findings indicate that smoking is associated with a loss of club cells, increased numbers of goblet cells, and substantial GCH airway remodeling.

### The bronchial airways of smokers contain a novel subpopulation of peri-goblet epithelial cells

We sought to establish the identity of cluster C-9, which was strongly enriched with current smoker cells and did not express established cell type marker genes (e.g. KRT5, FOXJ1, SCGB1A1, MUC5AC) (Figure 2c). C-9 cells expressed high levels of gene set GS-12, which contains the luminal cytokeratin KRT8 (Figure 5a). Additional cytokeratin genes were also present in GS-12, such as KRT13 and KRT19, as well as anti-oxidant genes, like TXN and GPX1 (Figure 5a). Cluster C-9 also expressed gene set GS-16, which was detected at low levels in MUC5AC^+^ cells (C-3) and contained the xenobiotic metabolism gene, CYP1B1 (Figure 5a). Furthermore, high expression of gene set GS-15 was detected in both C-9 cells and MUC5AC^+^ cells (C-3) (Figure 5a-c), suggesting that this cluster may have a functional relationship with goblet cells. GS-15 contains several genes previously reported to be persistently up-regulated post-smoking cessation (e.g. CEACAM5, CEACAM6, UPK1B)^18^, one of which has been explicitly linked to lung squamous cell carcinoma (SCC) and premalignancy (CEACAM5)^37^.

**Figure 5.**
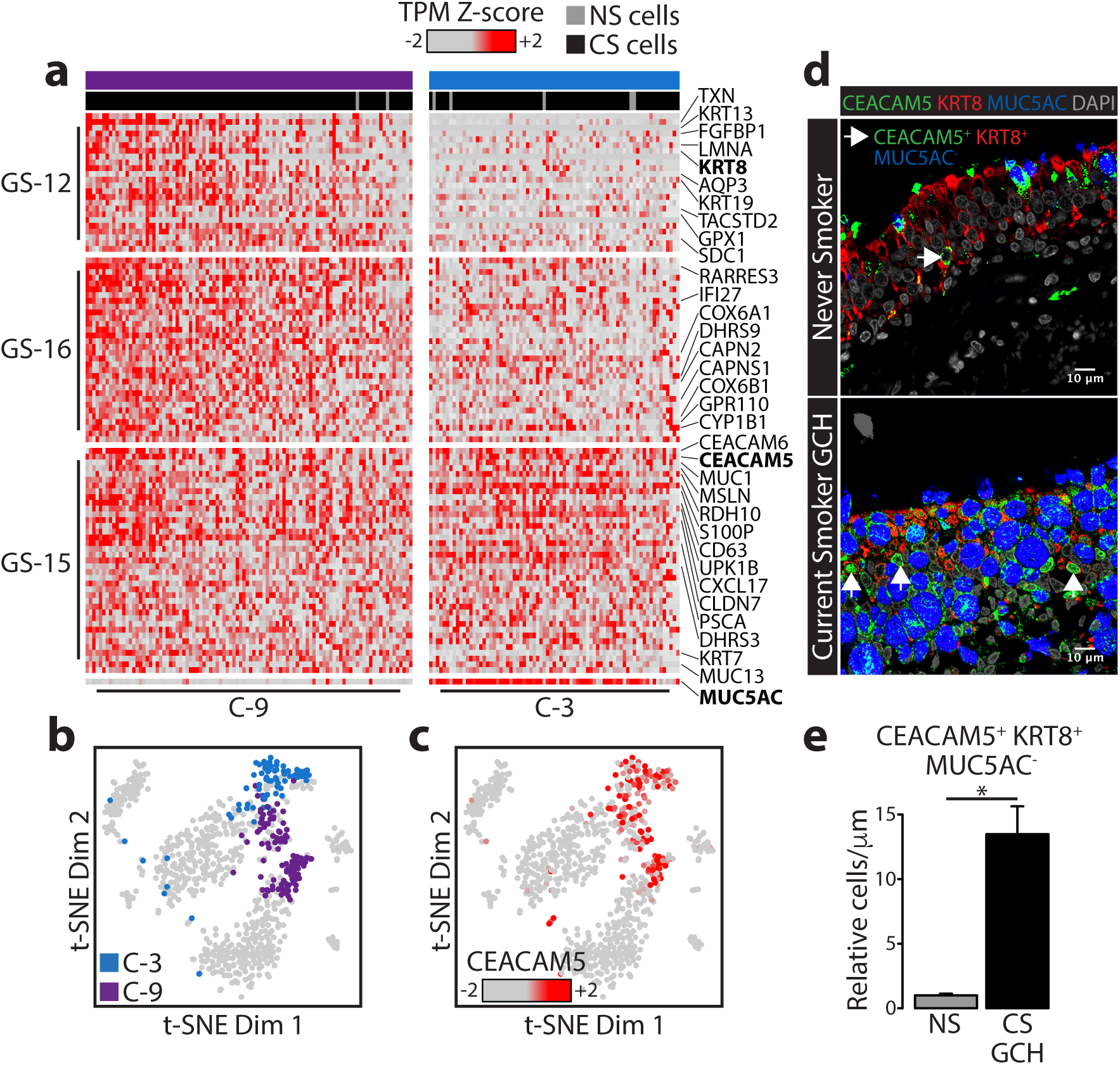
A novel subpopulation of peri-goblet cells was identified in the airways of smokers. (**a**) Expression of gene sets GS-12, GS-16, GS-15, and MUC5AC in clusters C-3 and C-9 was visualized by heatmap (z-normalized TPM values). (**b**) t-SNE was used to visualize cluster C-3 and C-9 cells as well as (**c**) CEACAM5 expression (z- normalized TPM values) across all cells. (**d**) Bronchial tissue procured from an independent cohort of never and current smokers (UCL Cohort, **Supplementary Table** was immunostained for CEACAM5, KRT8, and MUC5AC. Representative images of never smoker tissue (left), morphologically normal current smoker tissue (middle), and current smoker goblet cell hyperplasia (GCH) (right) were displayed. Changes in tissue length (μm)-normalized numbers of (**e**) CEACAM5^+^ KRT8^+^ MUC5AC^−^ cells in current smoker morphologically normal and GCH tissue were assessed relative to never smokers by WRS test.

To validate the presence of cluster C-9 cells in the airways of current smokers, bronchial tissue procured from a second independent cohort of never- and current smokers (UCL Cohort, **Supplementary Table 3**) was immunostained for KRT8, MUC5AC (goblet cells), and Ac-α-Tub (ciliated cells). KRT8^+^ MUC5AC^−^ Ac-α-Tub^−^ cells that were morphologically distinct from goblet- and ciliated cells were detected in significantly higher numbers in GCH regions of current smokers, relative to never smokers (**Supplementary Figure 15**). To confirm that there was functional overlap between goblet cells and this subpopulation of KRT8^+^ MUC5AC^−^ Ac-α-Tub^−^ cells, bronchial tissue (UCL Cohort, **Supplementary Table 3**) was immunostained for CEACAM5, in addition to KRT8 and MUC5AC. Increased numbers of CEACAM5^+^ KRT8^+^ MUC5AC^−^ cells were detected in GCH regions of current smokers, relative to never smokers (*p* = 0.004) (Figure 5d-e). Within current smoker GCH tissue regions, CEACAM5^+^ KRT8^+^ MUC5AC^−^ cells were typically found in close proximity to goblet cells (CEACAM5^+^ KRT8^+^ MUC5AC^+^) and were therefore named peri-goblet (PG) cells. CEACAM5 expression in goblet cells was phenotypically punctate and co-localized with MUC5AC in both never and current smokers (Figure 5d). In PG cells, however, CEACAM5 distinctively localized to the cell surface and nucleus (Figure 5d). Overall, this data indicates that PG cells are a novel, bronchial epithelial subpopulation associated with smoking-induced GCH.

## DISCUSSION

Previous transcriptomic studies have shown that smoking is associated with a robust bronchial gene expression signature^17-18^. Interrogation of bronchial tissue at single-cell resolution revealed that elements of this signature were derived from different cell subpopulations. Overall, we discovered smoking-associated phenotypes that included a metabolic response that localized to ciliated cells, a cell type shift that involved club cell loss and goblet cell expansion, and a previously uncharacterized subpopulation of peri-goblet (PG) epithelial cells present within regions of goblet cell hyperplasia.

We identified a gene set (GS-8) specifically expressed by smoker ciliated cells (C-5) that contains genes encoding families of enzymes, such as aldehyde dehydrogenases (e.g. ALDH3A1, ALDH1A3) and aldo-keto reductases (e.g. AKR1B10, AKR1C1), capable of breaking down tobacco smoke-derived chemical compounds, like toxic aldehydes (e.g. formaldehyde, acrolein) and ketones (e.g. acetone, methyl vinyl ketone)^8-9^. This finding suggests that ciliated cells exhibit a cell type-specific coping mechanism that may convey resistance to certain forms of smoking-induced toxicity. Links between this mechanism and previously reported smoking phenotypes, such as reduced ciliary length^15^, however, are unclear. This finding might also highlight a protective function with tissue-wide significance, in which the bronchial epithelium’s capacity for detoxification may be compromised if ciliated cells are lost due to injury or disease.

Several studies have reported that smoking is associated with increased mucous production and goblet cell hyperplasia in the bronchus^12-14,38-40^. Loss of club cells (SCGB1A1^+^) has been reported in smoker bronchioles^11-12^, but this is the first instance in which a similar observation has been made in the mainstem bronchus. We confirmed that goblet cell hyperplasia (GCH) is a regional phenomenon interspersed amongst morphologically normal (MN) tissue areas. The determinants of GCH prevalence are unclear, but it has been shown that cytokines (e.g. IL-13 and IL-4)^41-43^ and viral infection (e.g. Rhino virus and Poly(I:C))^44-45^ can increase MUC5AC expression and goblet cell abundance. The specific catalyst for GCH in response to smoke exposure is unknown, but reports of its co-occurrence with airway inflammation suggest that immunological interplay may be a factor^14^. Furthermore, there is evidence that both basal and club cells are capable of goblet cell differentiation^32,46^. However, the origins of newly formed goblet cells in the airways of smokers have not been explicitly described. Functional implications for goblet cell expansion and club cell loss are unclear, but a similar phenotype has been described in the airways of asthmatics, in which diminished mucosal fluidity, the formation of mucosal plugs, and impaired mucociliary clearance was observed^47-48^. Murine models have also shown that MUC5B loss is associated with impaired mucociliary clearance, airflow obstruction, and respiratory infection^49^.

Smoking-induced GCH was associated with the presence of a previously uncharacterized subpopulation of CEACAM5^+^ KRT8^+^ MUC5AC^−^ peri-goblet (PG) epithelial cells. The origins of PG cells are unclear, but a KRT8^+^ undifferentiated epithelial subpopulation derived from basal cells, referred to as “suprabasal”, has been described in murine models^46,50^. Suprabasal cells act as intermediate precursors to ciliated and secretory cells during basal cell differentiation under normal conditions^46^ and post-injury, as a repair mechanism^50^. However, the suprabasal phenomenon has not been characterized in the human bronchus and little is known regarding human intermediate epithelial subpopulations. Furthermore, the involvement of a KRT8^+^ intermediate state in club cell trans-differentiation^4,34^ has not been explored. Goblet cell differentiation required for the onset and maintenance of smoking-associated GCH might involve a pro-goblet precursor subpopulation, but the explicit role of PG cells in this context requires further investigation.

It has been reported that CEACAM5 expression is persistently up-regulated in the airways of former smokers, whereas genes specifically expressed by goblet cells, such as MUC5AC, SPDEF, and AGR2, return to normal, never smoker levels post-smoking cessation^18^. These findings suggest that goblet cell expansion in the airways of smokers is reversible, whereas the emergence of CEACAM5^+^ PG cells might have long-term implications. The functional consequences of the presence of PG cells are unclear, but irreversible alterations to bronchial epithelial composition might underlie chronic disease states. Although PG cells were identified in this study in the absence of established disease phenotypes, CEACAM5^+^ KRT5^+^ cells have been detected in bronchial premalignant lesions and lung SCC^37^. Therefore, investigation of mechanisms linking GCH-associated CEACAM5^+^ PG cells and premalignant lesion-associated CEACAM5^+^ KRT5^+^ cells might provide insight into smoking-induced conditions that drive lung carcinogenesis. Furthermore, characterization of all bronchial epithelial alterations indentified in smokers in this study will enable us to more precisely define the foundation upon which abnormalities arise during the early stages of smoking-associated lung diseases.

## METHODS

### Bronchial tissue collection for single-cell RNA sequencing

At Boston University Medical Center (BUMC), healthy, volunteer never smokers (n=6) and current smokers (n=6) underwent a bronchoscopy to obtain brushings from the right main-stem bronchus as described previously described^17-18^. Eligible volunteers included subjects who were 1) between the ages of 19 and 55; 2) did not use inhaled or intranasal medications; 3) did not have history of COPD, asthma, pulmonary fibrosis, or sarcoid, head & neck, or lung cancer; 4) did not use marijuana; 5) did not have a respiratory infection within the past 6 weeks; and 6) did not use other tobacco products (i.e. pipe, cigar, chewing). Spirometry was performed to assess lung function (e.g. FEV1/FVC). Exhaled carbon monoxide (Smokerlyzer Carbon Monoxide Monitor, Model EC-50; Bedfont Scientific Ltd) and urine cotinine (NicAlert; Confirm BioSciences) levels were measured to confirm smoking status. The Institutional Review Board approved the study and all subjects provided written informed consent.

### Single-Cell isolation by FACS

Bronchial brushings were treated with 0.25% Trypsin/EDTA for 20 minutes and stained for sorting using a BD FACSAria II. Gating based on forward scatter height vs. forward scatter area (FSC-H vs. FSC-A) was applied to sort only singlet events (**Supplementary Figure 1a**). Dead cells (LIVE/DEAD Fixable Aqua Dead Cell Stain, ThermoFisher; L34957) and red blood cells expressing GYPA/B (**Supplementary Figure 1b**) on their surface (APC anti-CD235ab; BioLegend; 306607) were stained and excluded. ALCAM^+^ epithelial cells (PE anti-CD166; BioLegend; 343903) and CD45^+^ white blood cells (APC-Cy7 anti-CD45; BD; 561863) were stained (**Supplementary Figure 1c**) and sorted into 96-well PCR plates containing lysis buffer (0.2% Triton-X 100, 2.5% RNaseOUT (ThermoFisher; 10777019)) compatible with downstream RNA library preparation. In each 96-well PCR plate for each subject, we sorted 84 ALCAM^+^ cells, 11 CD45^+^ cells, and maintained one empty well as a negative control. The plates were frozen on dry ice, and stored at −80 °C until preparation for sequencing.

### Single-cell RNA sequencing

Massively parallel single-cell RNA-sequencing of human bronchial airway cells was performed using a modified version of the CEL-Seq RNA library preparation protocol^20^. For each of the 12 recruited donors, one frozen 96-well PCR plate containing sorted cells was thawed on ice and RNA was directly reverse transcribed (ThermoFisher; AM1751) from whole cell lysate using primers composed of an anchored poly(dT), the 5’ Illumina adaptor sequence, a 6-nucleotide well-specific barcode, a 5-nucleotide unique molecular identifier (UMI), and a T7 RNA polymerase promoter. All primer sequences were listed in **Extended Supplementary Table 1**. Samples were additionally supplemented with ERCC RNA Spike-In mix (ThermoFisher; 4456740) (1:1,000,000 dilution) for quality control. cDNA generated from each of the 96 cells per plate was pooled, subjected to second strand synthesis (ThermoFisher; AM1751) and amplified by *in vitro* transcription (ThermoFisher; AM1751). Amplified RNA was chemically fragmented (New England BioLabs; E6150) and ligated to the Illumina RNA 3’ adapter (Illumina RS-200-0012). Samples were again reverse transcribed using a 3’ adaptor-specific primer and amplified using indexed Illumina RNA PCR primers (Illumina RS- 200-0012). In total, 1152 samples (1008 epithelial cells, 132 white blood cells, 12 negative controls) were sequenced on an Illumina HiSeq 2500 in rapid mode, generating paired-end reads (15 nucleotides for Read 1, 7 nucleotides for index, 52 nucleotides for Read 2).

### Data preprocessing

Illumina’s bcl2fastq2 software (v2.19.1) was used to de-multiplex the sequencing output to 12 plate-level FASTQ files (1 per 96-well plate). A python-based pipeline (https://github.com/yanailab/CEL-Seq-pipeline) was utilized to: 1) de-multiplex each plate-level FASTQ file to 96 cell-level FASTQ files, trim 52 nucleotide reads to 35 nucleotides, and append UMI information from read 1 (R1) to the header of read 2 (R2); perform genomic alignment of R2 with Bowtie2 (v2.2.2) using a concatenated hg19/ERCC reference assembly; and 3) convert aligned reads to gene-level counts using a modified version of the HTSeq (v0.5.4p1) python library that identifies reads aligning to the same location with identical UMIs and reduces them to a single count. One UMI-corrected count was then referred to as a “transcript”. The pipeline was configured with the following settings: alignment quality (min_bc_quality) = 10, barcode length (bc_length) = 6, UMI length (umi_length) = 5, cut_length = 35.

### Data quality control

The quality of each cell was assessed by examining the total number of reads, total reads aligned to hg19, total reads aligning to genes (pre-UMI correction), total transcript counts, and total genes with at least one detected transcript. Cells were excluded from downstream analyses if the total number of transcripts were not two-fold greater than the total background-level transcripts detected in the empty well negative control on each plate (**Supplementary Figure 3**). Cells were also excluded from downstream analyses if there was a weak Pearson correlation (*r* < 0.7) between detected ERCC RNA Spike-in transcript counts (Log_10_) and ERCC input concentration (Log_10_) (amol/mL) (**Supplementary Figure 3**). All non-protein-coding genes and genes with less than 2 transcript counts in 5 cells were removed from the dataset. The remaining 7680 genes measured across 796 cells were used for subsequent analyses.

### Latent Dirichlet allocation (LDA) implementation and model optimization

Latent Dirichlet Allocation (LDA) from the topicmodels R package (v0.2-6) was used to generate probabilistic representations of cell clusters and gene sets present in the dataset, referred to as Cell-States and Gene-States. The input for the Cell-State model required a counts data matrix where cells were columns and genes were rows, whereas for the Gene-State model, the same matrix was transposed (ie. genes were columns and cells were rows). Models were fit using the Variational Expectation-Maximization (VEM) algorithm with the following parameters: nstart = 5, seed = 12345, estimate.alpha = TRUE, estimate.beta = TRUE. The given parameter “*k*” determined the number of Cell-States and Gene-States to be estimated by the model. The optimal value of k was determined by 5-fold cross validation and evaluation of model perplexity. For the Gene-State model, cells were randomly partitioned into “training” (80%) and “test” (20%) sets, whereas for the Cell-State model, genes were randomly partitioned into “training” (80%) and “test” (20%) sets. Models were then fit to the training set and perplexity was estimated to evaluate model fit for the held-out test set. Fifty iterations of this process were performed for *k* = 2-50, mean perplexity was calculated at each *k*, and the minimum mean perplexity was selected as the optimal value of *k* (ie. *k.opt*), which was *k* = 13 for the Cell-State model and *k* = 19 for the Gene-State model (**Supplementary Figure 6**).

### Gene set (GS) and cell cluster (C) assignments

Negative binomial generalized linear models were built using the MASS R package (v7.3-45) for each Gene-State (n=19) and each Cell-State (n=13), in which States were treated as inferred, independent variables and genes or cells, respectively, were treated as dependent variables. A cell was assigned to a Cell-State if a significant association (FDR *q* < 0.05) was observed with positive directionality (regression coefficient > 1). A Similarly, a gene was assigned to a Gene-State if a significant, positive association was observed (FDR *q* < 1 x 10-5, regression coefficient > 1). If multiple State associations were observed for a given gene or cell, assignment was determined based on the strongest State association (ie. minimum FDR *q*). Additional metrics for gene set and cluster assignment include *State Specificity* and *State Similarity*. LDA (see Methods: Latent Dirichlet allocation (LDA) implementation and model optimization) also assigned a probability to each gene (or cell) for each Gene-State (or Cell-State) and *State Specificity* was calculated by dividing that probability by the sum of probabilities across all Gene-States (or Cell-States). A minimum *State Specificity* of 0.1 was required for gene or cell assignment. *State Similarity* was calculated by assessing the cosine (θ) similarity between each Gene-State and relative expression of each gene (gene counts divided by total counts for each cell). A minimum *State Similarity* of 0.4 was required for gene assignment. All downstream analyses used the 785 cells that fit the criteria for Cell-State assignment and 676 genes that fit the criteria for Gene-State assignment. Statistical modeling results, *State Specificity*, and *State Similarity* values for all genes, regardless of assignment status, were included in **Extended Supplementary Table 2**.

### Data visualization by heatmap and t-SNE

Prior to heatmap visualization, transcript counts were transformed to z-normalized transcripts per million (TPM). Genes (top-to-bottom) and cells (left-to-right) were ordered according to the strength of statistical association (FDR *q*) with respective assigned Gene-States and Cell-States. The tsne R package v0.1-3 was used for dimensionality reduction by t-distributed stochastic neighbor embedding (t-SNE). Modified parameters include: k = 2, seed = 1234. Input for t-SNE was z-normalized transcripts per million (TPM) values across genes with at least 3 transcript counts in 3 cells (n = 4,914 genes). Gene expression overlay onto t-SNE visualization was also performed using z-normalized TPM values.

### Functional annotation

The enrichR R package (v0.0.0.9000) was used as an interface for the web-based functional annotation tool, Enrichr, to identify Gene Ontology terms from the GO Biological Process 2015 library significantly associated with each gene set^51-52^. Functional annotation results were listed in **Extended Supplementary Table 3**.

### Microarray data processing

Raw CEL files obtained from the Gene Expression Omnibus (GEO) for Series GSE7895 were normalized to produce gene-level expression values using the implementation of the Robust Multiarray Average (RMA) in the affy R package (v1.36.1) and an Entrez Gene-specific probeset mapping (17.0.0) from the Molecular and Behavioral Neuroscience Institute (Brainarray) at the University of Michigan (http://brainarray.mbni.med.umich.edu/>).

### Comparative analysis of scRNA-Seq and microarray data

Bronchial brushings were reconstructed *in silico* from the single-cell data by taking the sum of all transcript counts for each gene across all cells procured from each donor. Negative binomial generalized linear models were built using the MASS R package (v7.3-45), modeling transcript counts as a function of smoking status (FDR *q* < 0.05: n=593 genes). In parallel, using never and current smoker bulk bronchial brushing microarray data (GEO series GSE7895), linear models were built using the stats R package (R v3.2.0), modeling gene-level expression values as a function of smoking status (FDR *q* < 0.05: n=689 genes). The correlation between test statistics generated from both models was then measured to compare differential expression results (**Supplementary Figure 4a**). Using the overlap amongst smoking-associated genes identified in both models (n=155 genes), correlations (Spearman) amongst *in silico* bronchial brushings and bulk bronchial brushings were examined (**Supplementary Figure 4b**).

### Gene set expression analysis in microarray data

Using published microarray data generated from bulk bronchial brushings procured from never and current smokers (GEO Series GSE7895), RMA-transformed values for each gene were z-normalized. MetaGene values were then generated by computing the mean z-score across all genes in each gene set (GS-1 to 19) for each sample. For each MetaGene, differential expression between never and current smoker samples was assessed using the Wilcoxon Rank Sum (WRS) test. For each smoking-associated MetaGene (*p* < 0.05), if the mean current smoker value was greater than- or less than- the mean never smoker value, the gene set was considered to be up- or down-regulated in current smokers, respectively.

### Cell type assessment for cell clusters

TPM values for cell type marker genes (KRT5, FOXJ1, SCGB1A1, MUC5AC, and CD45) were z-normalized across all cells. Cluster-specific mean expression was designated: high (pink) if expression exceeded one standard deviation above the mean value across all cells; medium (white) if expression exceeded one half of a standard deviation above the mean value across all cells; low (light grey) if expression exceeded the mean value across all cells. If cluster-specific mean expression was designated high, medium, or low for KRT5, FOXJ1, SCGB1A1, MUC5AC, or CD45 (PTPRC), that cluster was assigned the cell type of basal, ciliated, club, goblet, or white blood cell, respectively. Cluster-specific mean expression below the mean value across all cells indicated that a given cluster did not express a given marker gene (dark grey).

### Gene set expression analysis in cell clusters

Transcript counts were transformed to z-normalized transcripts per million (TPM). MetaGene values were then generated by computing the mean z-score across all genes in each gene set (GS-1 to 19) for each cell. Cluster-specific MetaGene expression was designated: high (pink) if mean expression exceeded one standard deviation above the mean value across all cells; medium (white) if mean expression exceeded one half of a standard deviation above the mean value across all cells; low (light grey) if mean expression exceeded the mean value across all cells. Cluster-specific mean expression below the mean value across all cells indicated that a given cluster did not express a given gene set (dark grey).

### Bronchial tissue collection for immunostaining

Bronchial tissue was collected from patients undergoing lung resection. All specimens were procured at least 5 cm from bronchial sites affected by disease diagnoses and analyses indicated that tissue was histologically normal. The UMCG Cohort (**Supplementary Table 2**) included specimens analyzed in collaboration with University Medical Center Groningen (UMCG) collected from 4 never smokers and 4 current smokers. Specimens were obtained from the tissue bank in the UMCG Department of Pathology. The study protocol was consistent with the Research Code of the University Medical Center Groningen and Dutch national ethical and professional guidelines (“Code of conduct; Dutch federation of biomedical scientific societies”; http://www.federa.org). The UCL cohort (**Supplementary Table 3**) included specimens analyzed in collaboration with University College London collected from 5 never smokers and 5 current smokers. Ethical approval was sought and obtained from University College London (UCL) Hospital Research Ethics Committee (REC reference 06/Q0505/12). This study was carried out in accordance with the Declaration of Helsinki (2000) of the World Medical Association.

### Immunofluorescence

Formalin-fixed paraffin-embedded lung sections were cut at 4 μm, tissue was probed with primary antibodies (listed below) and secondary antibodies with fluorescent conjugates (Invitrogen Alexa Fluor 488, 594, 647), and nuclear staining was performed with DAPI (ThermoFisher; R37606). Immunostaining was performed using the following primary antibodies: mouse anti-Acetylated-a-Tubulin (Ac-a-Tub) (Sigma; T6793) (Sigma), rabbit anti-Acetylated-α-Tubulin (Ac-α-Tub) (Enzo Life Sciences; BML SA4592), rabbit anti-AKR1B10 (Sigma; HPA020280), rabbit anti-CEACAM5 (Abcam; ab131070), chicken anti-KRT5 (BioLegend; 905-901), rat anti-KRT8 (Developmental Studies Hybridoma Bank, University of Iowa; TROMA-I), mouse anti-MUC5AC (Abcam; ab3649), rabbit anti-MUC5B (Sigma; HPA008246). Imaging of staining panels analyzed in collaboration with investigators at University Medical Center Groningen (**Supplementary Table 2**) (e.g. AKR1B10 / Ac-α-Tub / KRT8, Figure 3c; AKR1B10 / MUC5AC / KRT8, **Supplementary Figure 13**; MUC5B / MUC5AC, Figure 4b; MUC5B / MUC5AC / KRT5, **Supplementary Figure 14**) was performed using a Carl Zeiss LSM 710 NLO confocal microscope at 63x objective magnification at the Boston University School of Medicine Multiphoton Microscope Core Facility. Imaging of staining panels analyzed in collaboration with investigators at University College London (**Supplementary Table 3**) (e.g. CEACAM5 / KRT8 / MUC5AC, Figure 5d; KRT8 / MUC5AC / Ac-α-Tub, **Supplementary Figure 15**) was performed using a Leica TCS Tandem confocal microscope at 63x objective magnification.

### Imaging analysis

All imaging data was analyzed using ImageJ Fiji software. For each image, cells were counted relative to the measured length of the epithelium in microns (cells/μm). Mean cell counts per micron (cells/μm) was then calculated for never smokers (treated as the control) and individual values for each image from never and current smokers were calculated relative to the never smoker mean (ie. relative cells/μm). We analyzed three images for each donor and assessed smoking-associated changes using the Wilcoxon rank-sum test. For panels in which MUC5AC was stained, current smoker tissue was assigned phenotypic status of either “morphologically normal” (MN) or “goblet cell hyperplasia” (GCH), based on qualitative assessment of goblet cell density and stratification. For each current smoker, three images of each status were analyzed.

## Supporting information

## ACKNOWLEDGEMENTS

We thank Dr. Itai Yanai at New York University School of Medicine for assistance with the CEL-Seq protocol; Dr. Brigitte Gomperts at University of California, Los Angeles for help with tissue dissociation and FACS; and Dr. Xaralabos Varelas at Boston University School of Medicine for providing fluorescence microscopy support.

This study was supported by funding from the Department of Defense grant W81XWH- 14-1-0234 (J.B.). J.B. and J.C. were supported by the LUNGevity Career Development Award. G.D. was supported by the NIH T32 training grant HL007035. S.J. is a Wellcome Trust Senior Fellow in Clinical Science and is supported by the Rosetrees Trust and UCLH Charitable Foundation. V.T. and S.J. are funded by the Roy Castle Lung Cancer Foundation. This work was partially undertaken at UCLH/UCL, which received funding from the UK Department of Health’s NIHR Biomedical Research Centre’s funding scheme (S.J.).

## AUTHOR CONTRIBUTIONS

Study conception and design (G.D., J.B., J.C., A.S., M.L.); collection of clinical samples (A.S., Y.G., R.T., Y.D.); sample processing (G.D., P.A.); library preparation and sequencing (G.D., G.L.); data analysis (G.D., J.B., J.C.); immunofluorescence (G.D., V.T., W.T., S.J., M.B., C.B., M.R.); manuscript writing (G.D., J.B., J.C.); manuscript editing (G.D., J.B., J.C., S.M., A.S., M.L., W.T., M.B., C.B.).

