## Supplementary material for "Characterizing smoking-induced transcriptional heterogeneity in the human bronchial epithelium at single-cell resolution"

| | Age | Sex | Pack years | FEV1/FVC | Cotinine (ng/mL) | CO (ppm) |
|---|---|---|---|---|---|---|
| Never Smoker 1 | 43 | Female | 0 | 88 | 10-30 | 3 |
| Never Smoker 2 | 34 | Male | 0 | 73 | 10-30 | 4 |
| Never Smoker 3 | 26 | Female | 0 | 83 | 10-30 | 3 |
| Never Smoker 4 | 25 | Male | 0 | 81 | 10-30 | 4 |
| Never Smoker 5 | 25 | Male | 0 | 83 | 10-30 | 3 |
| Never Smoker 6 | 24 | Female | 0 | 88 | 10-30 | 4 |
| Current Smoker 1 | 35 | Male | 11 | 84 | 1000 | 17 |
| Current Smoker 2 | 49 | Female | 14 | 81 | 1000 | 9 |
| Current Smoker 3 | 51 | Male | 13 | 78 | 1000 | 16 |
| Current Smoker 4 | 27 | Female | 9 | 89 | 1000 | 19 |
| Current Smoker 5 | 50 | Male | 31 | 83 | 1000 | 9 |
| Current Smoker 6 | 44 | Female | 14 | 81 | 1000 | 13 |

**Supplementary Table 1. Bronchial brushings were procured from 6 never smokers and 6 current smokers.** Never and current smoker donors were healthy volunteers recruited at Boston University Medical Center (BUMC). In addition to bronchoscopy and procurement of bronchial brushing, spirometry was performed to assess lung function by measuring forced expiratory volume in one second (FEV1) relative to forced vital capacity (FVC). Exhaled carbon monoxide (CO) and urine cotinine levels were also measured to confirm smoking status.

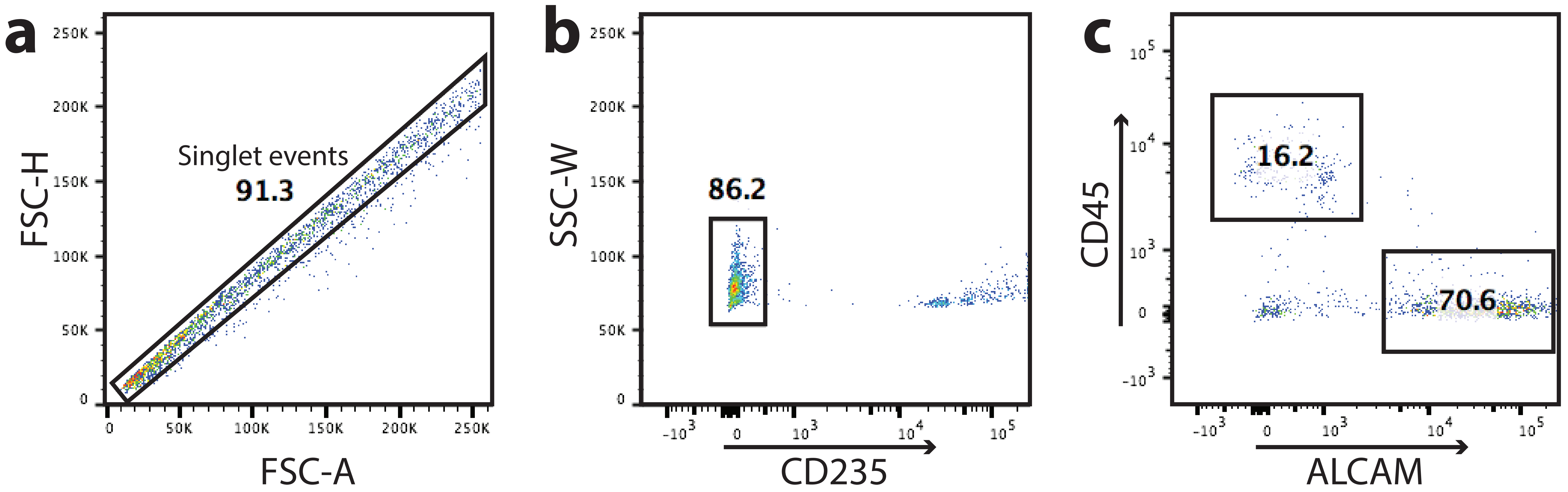


**Supplementary Figure 1. Single bronchial cells were isolated by FACS.** (**a**) Gating based on forward scatter height vs. forward scatter area (FSC-H vs. FSC-A) was applied to sort only singlet events (single cells). (**b**) Red blood cells expressing GYPA/B on their surface were stained and excluded. (**c**) Epithelial cells expressing ALCAM on their surface and WBCs expressing CD45 on their surface were stained and sorted.

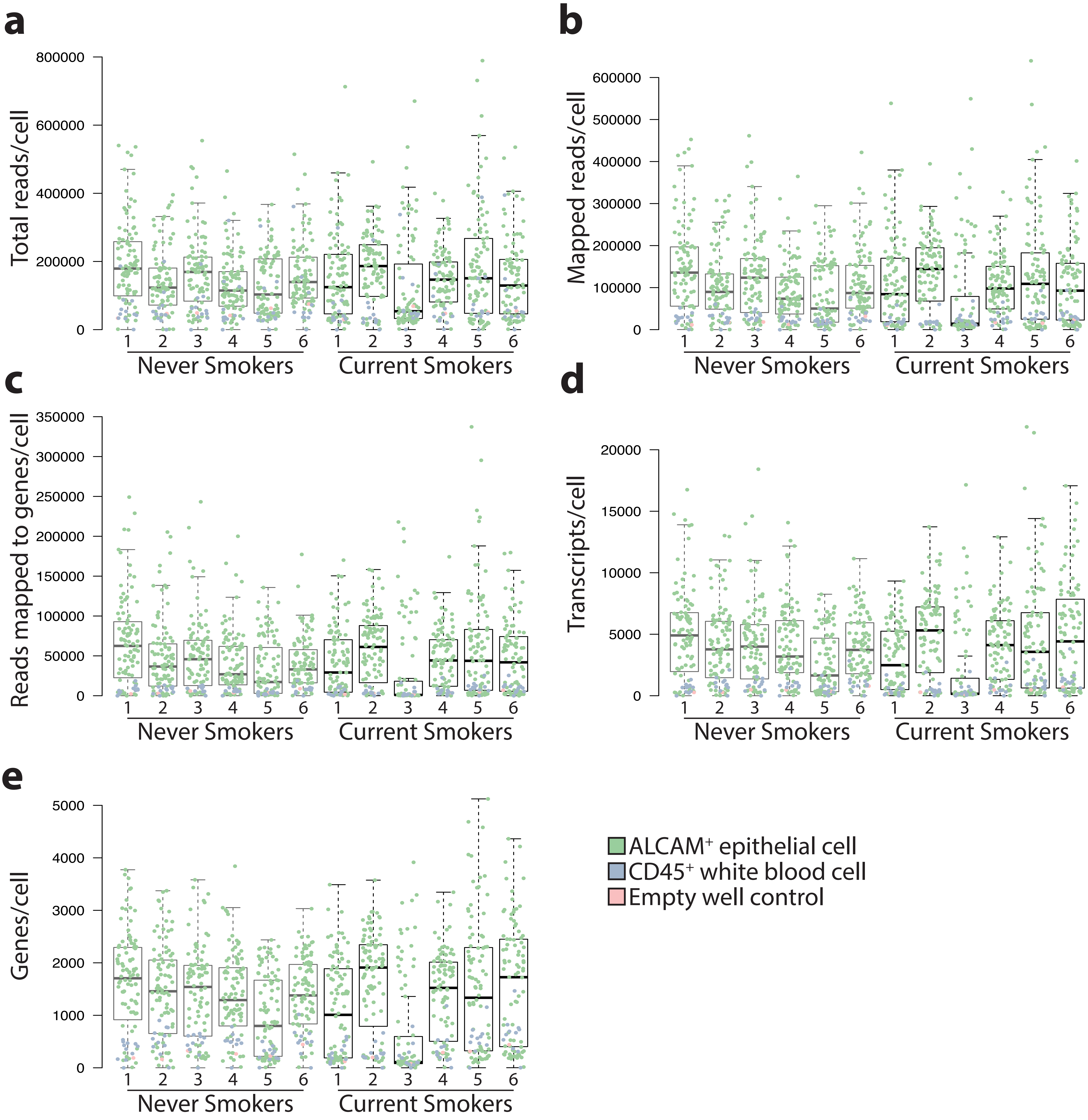


**Supplementary Figure 2. Single cell RNA-Seq data quality was evaluated for each donor.** For each cell from each donor, numbers of total reads (**a**), reads that mapped to hg19 (**b**), reads that mapped to genes (pre-UMI correction) (**c**), transcript counts (post-UMI correction) (**d**), and genes with at least one transcript count (**e**) were measured.

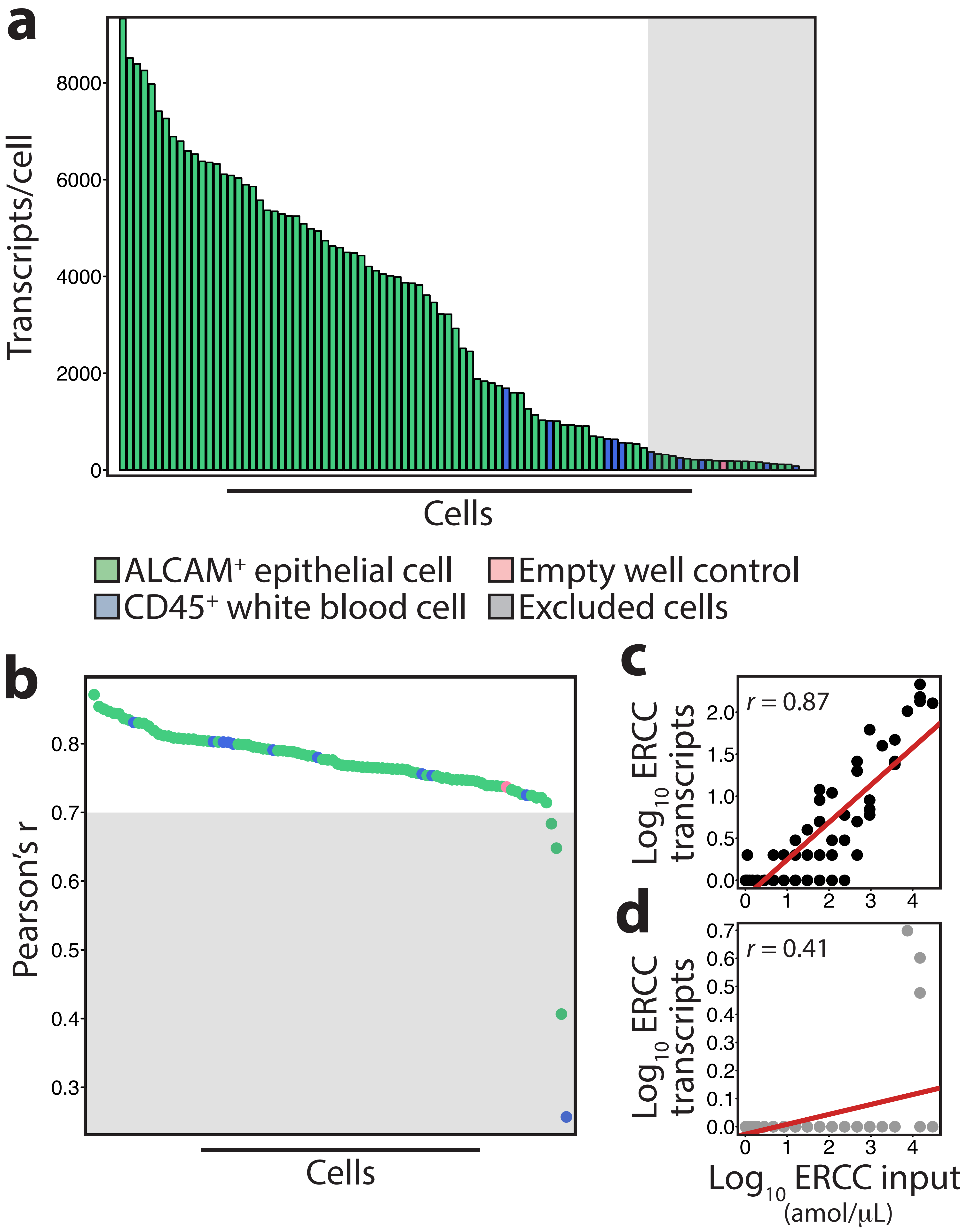


**Supplementary Figure 3. Low quality cells were excluded from downstream analyses.** Figures A-D are depicting data from a single donor: 84 epithelial cells, 11 WBCs, and 1 empty well negative control. (**a**) Total transcript counts for each cell were plotted and those with less than 2-fold the background-level counts detected in the empty well negative control (grey) were excluded from subsequent analyses. (**b**) Pearson correlation between detected ERCC RNA Spike-in transcript counts ($Log_{10}$) and ERCC input concentration ($Log_{10}$) (amol/ml) was plotted for each. Cells in which a weak correlation ($r < 0.7$) was observed were excluded from subsequent analyses. Examples of cells with (**c**) strong ($r = 0.87$) and (**D**) weak ($r = 0.41$) correlations between detected ERCC transcript counts and ERCC input were displayed.

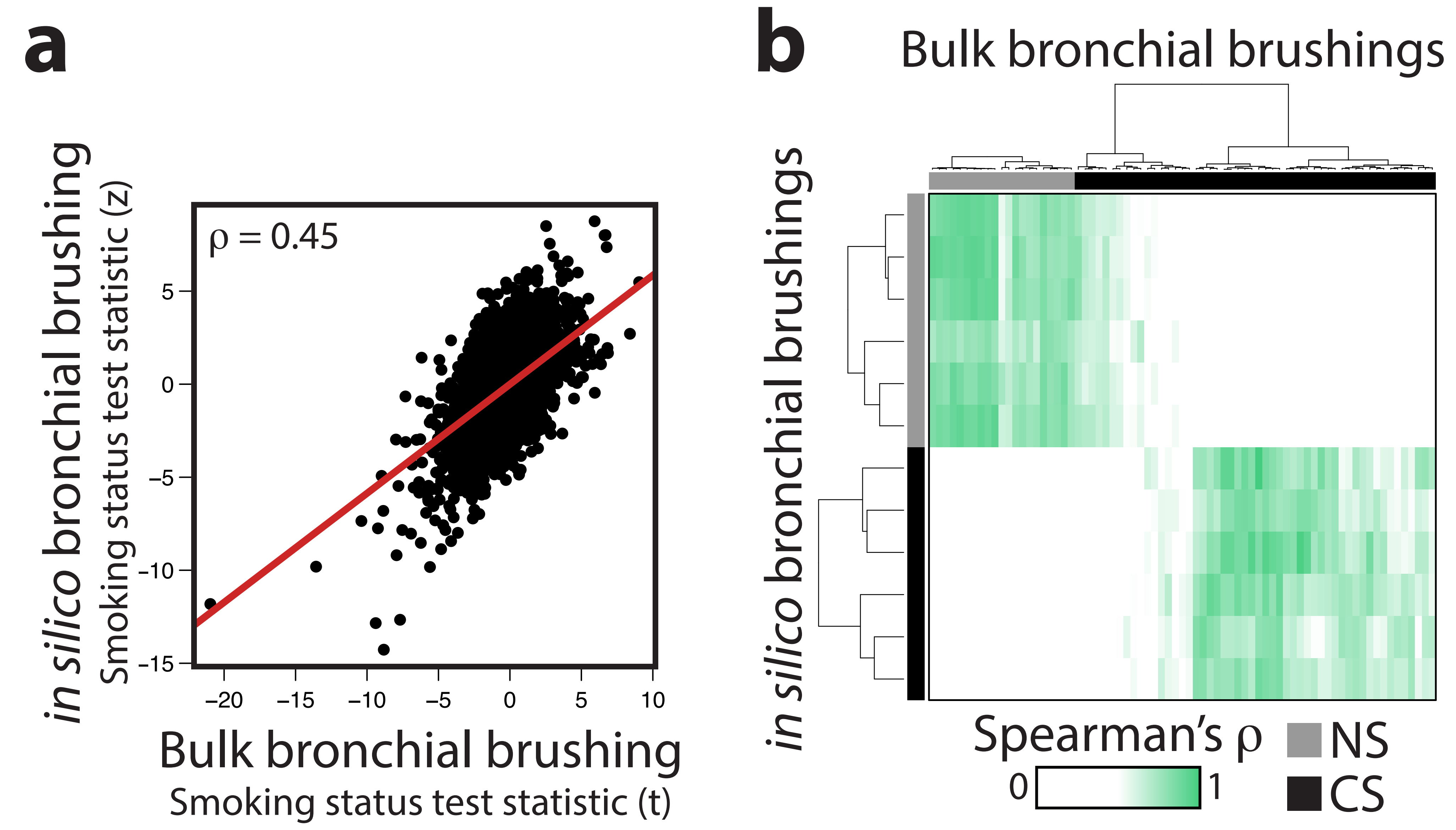


**Supplementary Figure 4. Bronchial brushings reconstructed *in silico* from single-cell data resemble data generated from bulk bronchial brushings.** For each donor, bronchial brushings were reconstructed in silico by combining data from all profiled cells isolated from each donor. (**a**) Differential expression between never and current smoker *in silico* bronchial brushings was compared to a similar analysis performed using published microarray data generated from bulk bronchial brushings. The correlation amongst test statistics generated from the two differential expression analyses was assessed. (**b**) Spearman correlations amongst in silico bronchial brushings and bulk bronchial brushing data were displayed by heatmap.

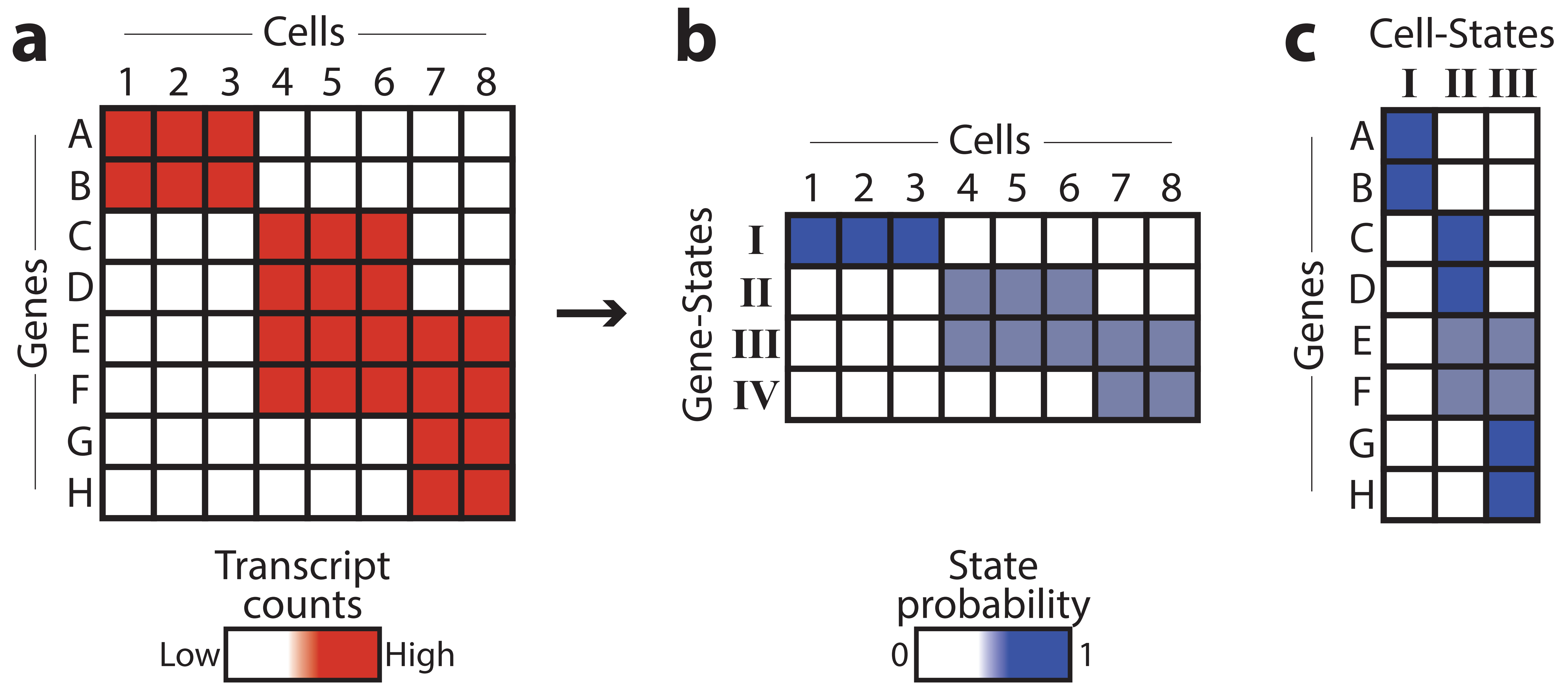


**Supplementary Figure 5. Latent Dirichlet Allocation (LDA) was used identify Cell-States and Gene-States.** (**a**) Model input for LDA was a data matrix containing single-cell transcript counts (rows = Genes, columns = Cells). (**b**) LDA was used to generate probabilistic representations of the types of genes present in the dataset, referred to as Gene-States (n=4: I-IV). (**c**) LDA was used to generate probabilistic representations of the types of cells present in the dataset, referred to as Cell-States (n=3: I-III).

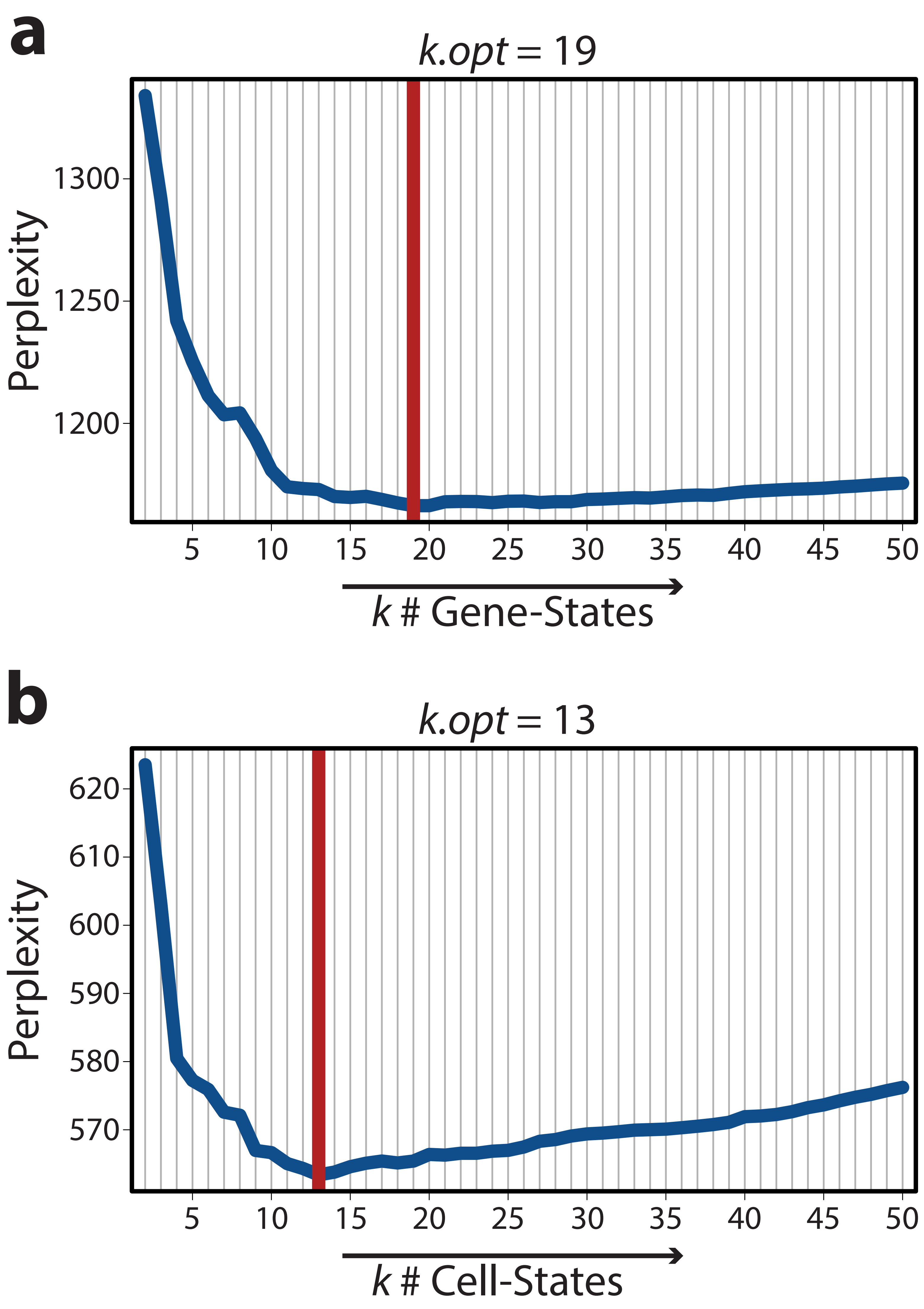


**Supplementary Figure 6. Gene-State and Cell-State models were optimized.** To determine the optimal number (k.opt) of Gene-States and Cell-States when performing LDA, model perplexity was evaluated for 50 iterations of 5-fold cross validation at k = 2-50. (**a**) For Gene-State models, mean perplexity was calculated across all iterations and plotted for each value of k. The minimum mean perplexity value was selected as k.opt (k = 19). (**b**) For Cell-State models, mean perplexity was calculated across all iterations and plotted for each value of k. The minimum mean perplexity value was selected as k.opt (k = 13).

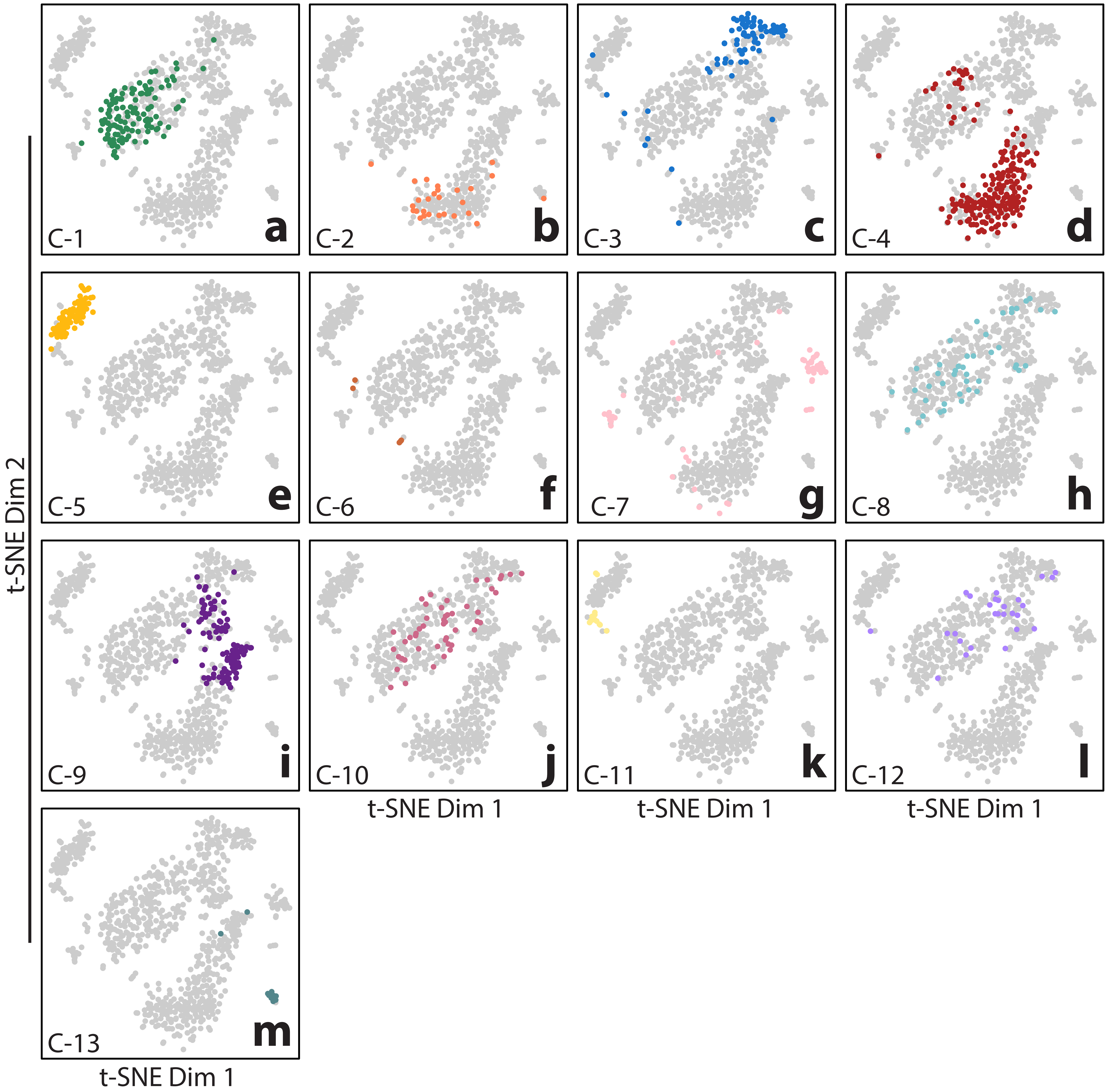


**Supplementary Figure 7. LDA was used to identify 13 cell clusters.** Each cell cluster (C-1 to 13) associated with an LDA-derived Cell-State was visualized by t-SNE (**a-m**).

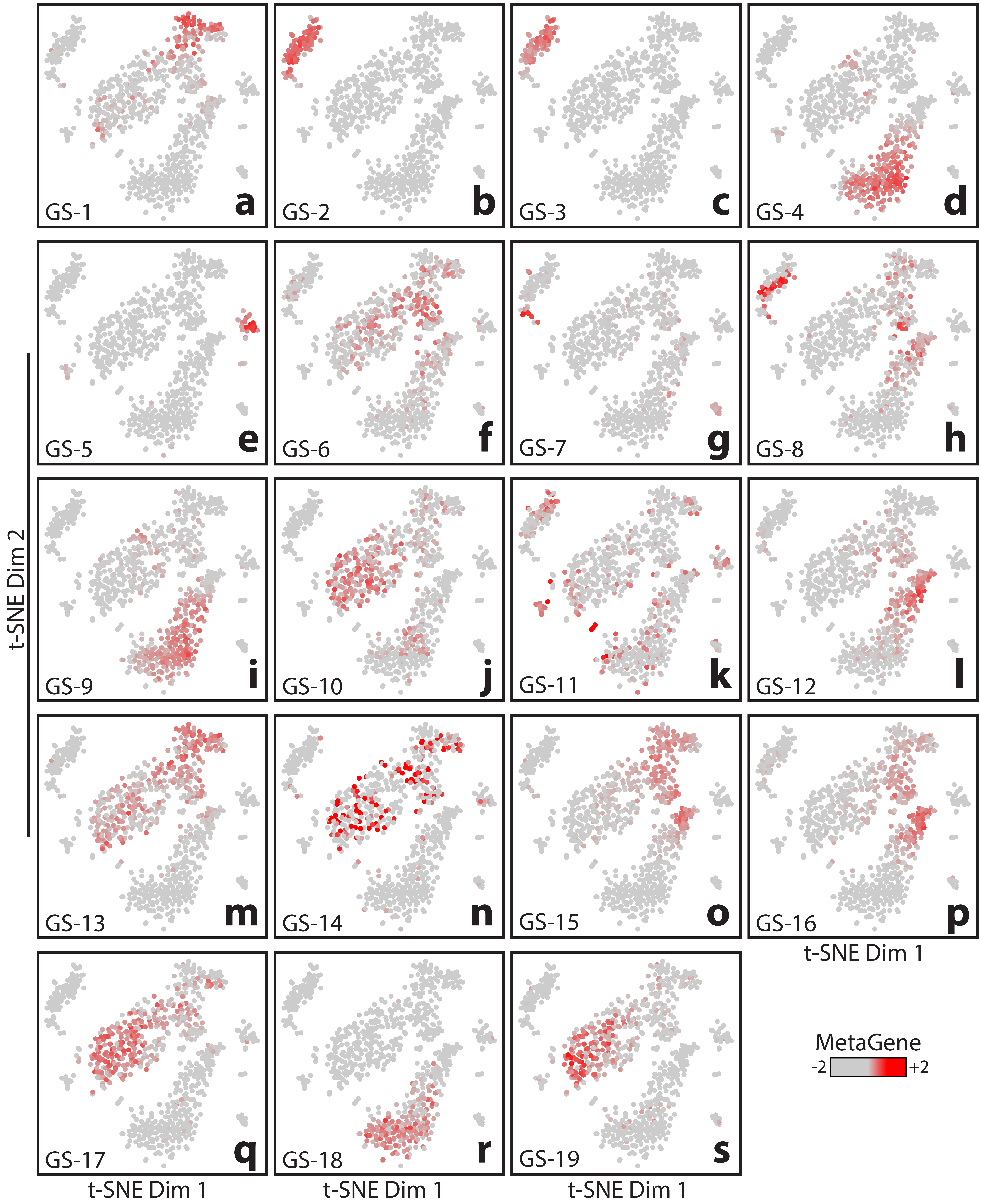


**Supplementary Figure 8. LDA was used to identify 19 gene sets.** A MetaGene was generated for each gene set (GS-1 to 19) associated with an LDA-derived Gene-State and expression values were visualized by t-SNE (**a-s**).

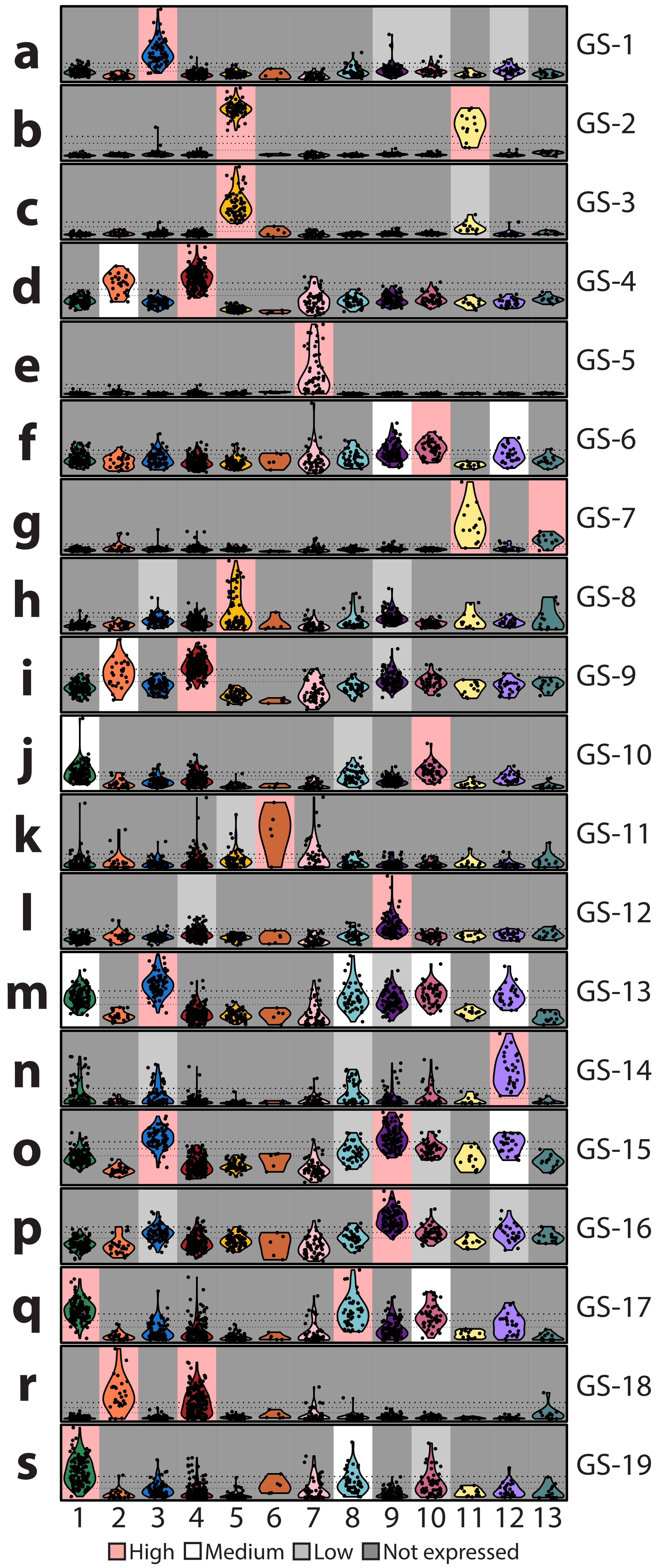


**Supplementary Figure 9. Gene set expression was assessed in each cluster.** A MetaGene was generated for each gene set (GS-1 to 19) and violin plots (**a-s**) were used to display the distribution of MetaGene expression values amongst cells in each cluster. Cluster-specific MetaGene expression was designated: high (pink), medium (white), low (light grey), or not expressed (dark grey). Horizontal dotted lines in each plot designated: mean expression across all cells (bottom), one half of a standard deviation above the mean value across all cells (middle), and one standard deviation above the mean value across all cells (top).

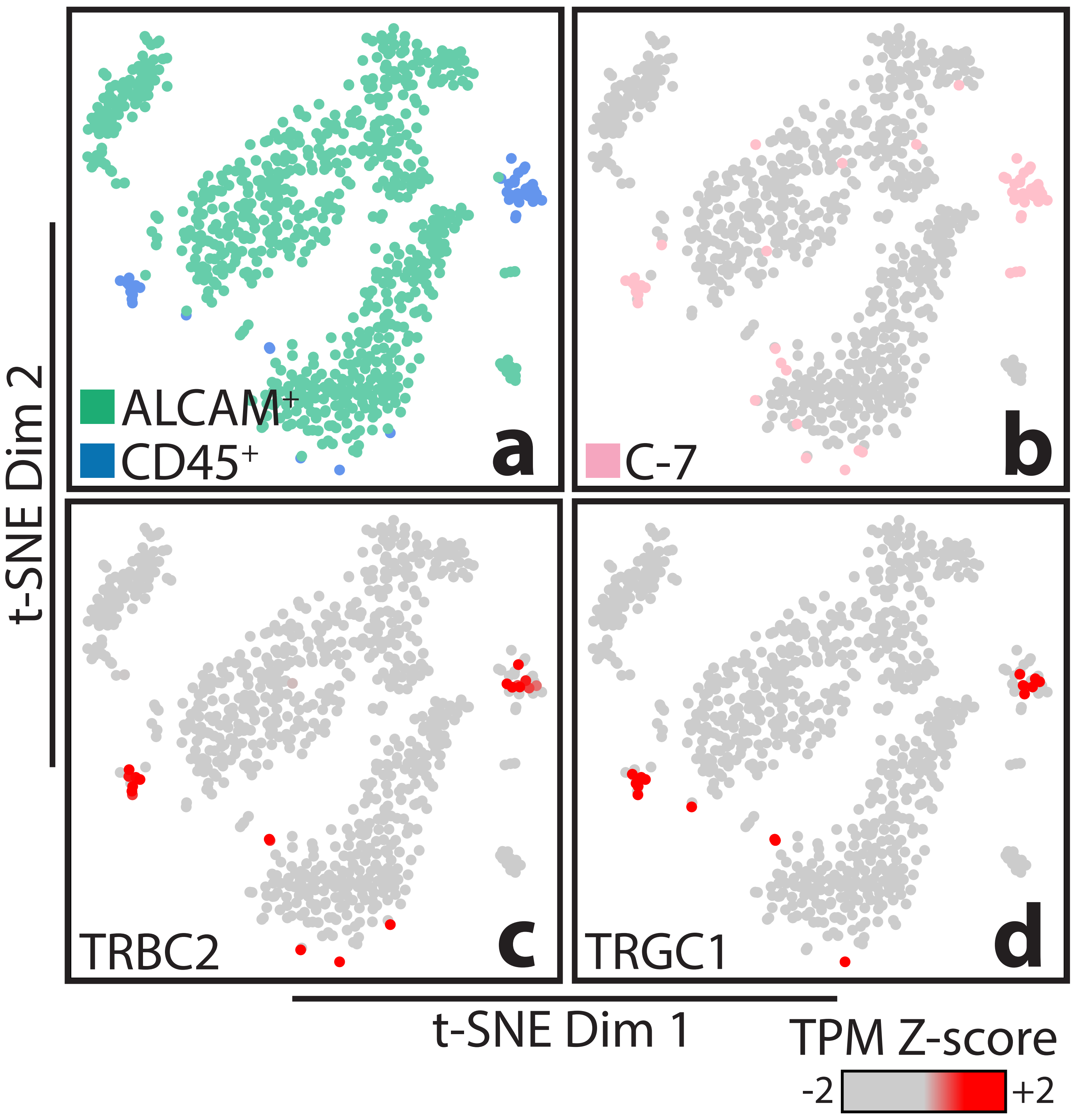


**Supplementary Figure 10. T cell receptor genes were detected in CD45+ cell cluster.** t-SNE was used to visualize (**a**) sorted ALCAM+ epithelial cells and CD45+ WBCs, (**b**) Cluster C-7 cells, and expression (z-normalized TPM values) of T cell receptor genes (**c**) TRBC2 and (**d**) TRGC1 across all cells.

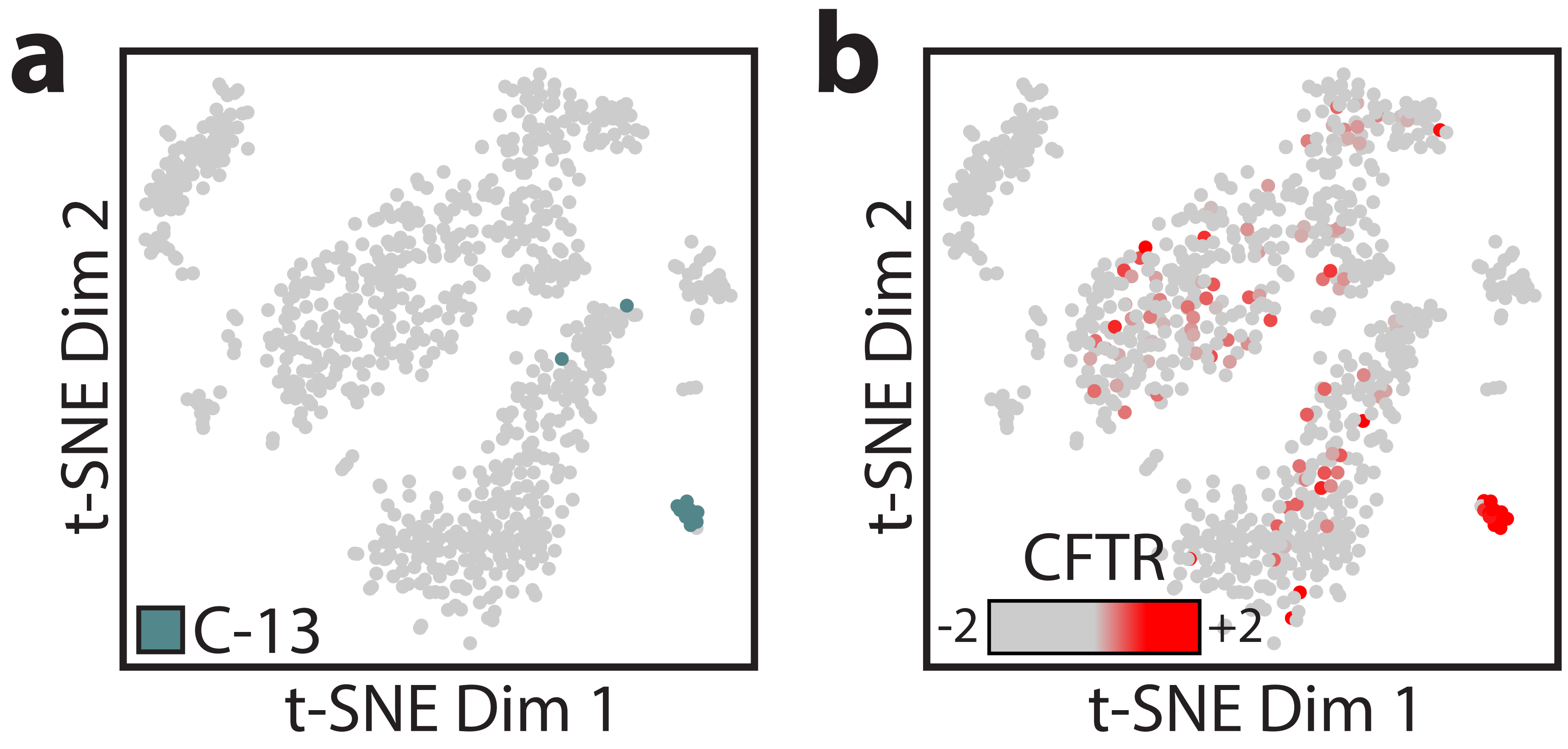


**Supplementary Figure 11. Cluster 13 cells expressed CFTR.** t-SNE was used to visualize (**a**) Clusters C-13 cells as well as (**b**) expression (z-normalized TPM values) of CFTR across all cells.

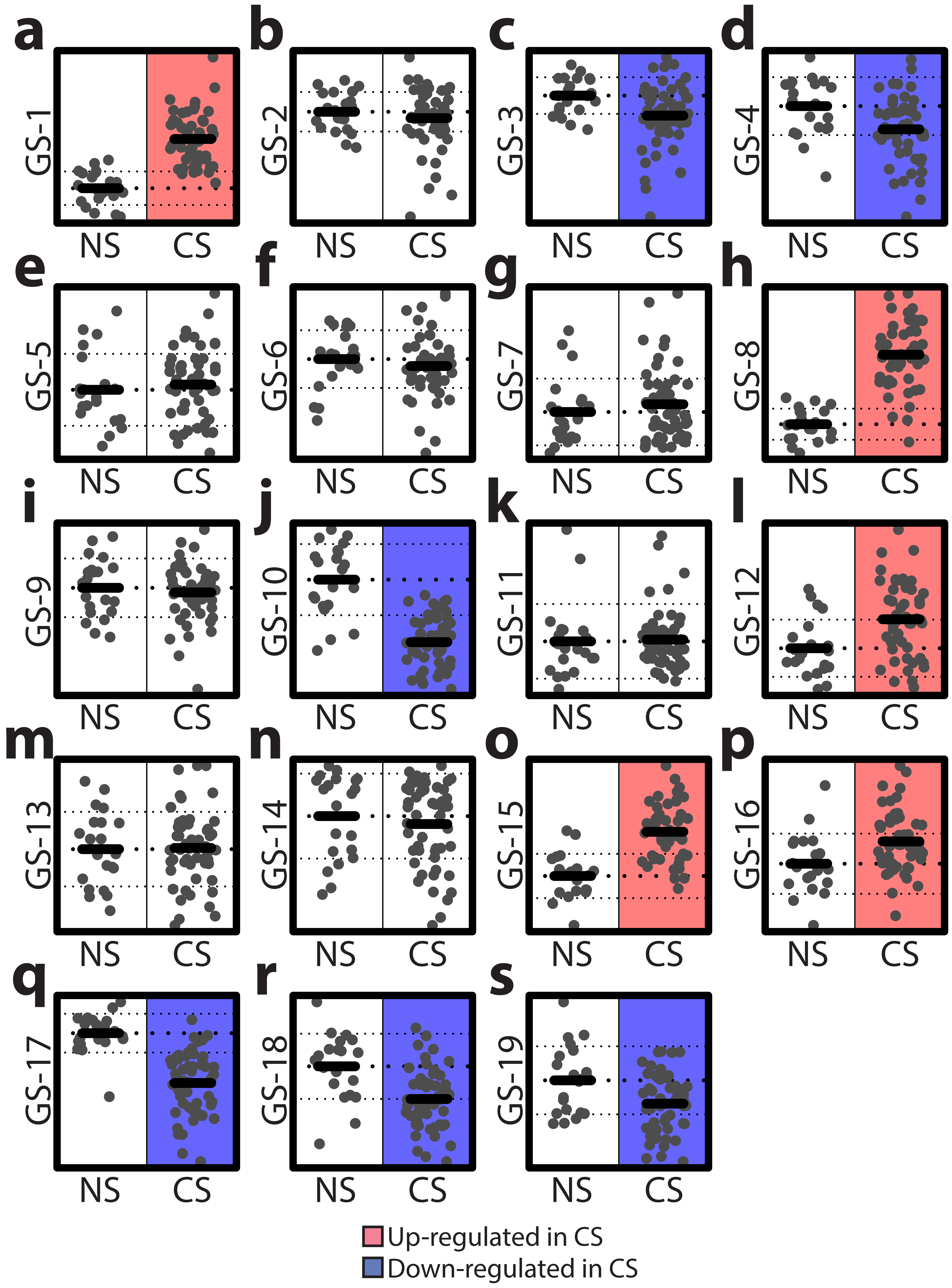


**Supplementary Figure 12. Smoking-associated differential expression of each gene set was assessed in published bulk bronchial brushing data.** Using published microarray data generated from bulk bronchial tissue procured from never (NS) and current (CS) smokers (GEO series GSE7895), MetaGene values were computed for each gene set (GS-1 to 19, **a-s**) and smoking-associated differential expression was assessed (WRS $p < 0.05$).

| | Age | Sex | Pack years | FEV1/FVC | Diagnosis |
|---|---|---|---|---|---|
| Never Smoker 1 | 45 | Male | 0 | 83 | IMT |
| Never Smoker 2 | 49 | Male | 0 | 78 | PC |
| Never Smoker 3 | 50 | Male | 0 | 78 | MET |
| Never Smoker 4 | 81 | Female | 0 | 73 | ADC |
| Current Smoker 1 | 45 | Female | 30 | 82 | LCC |
| Current Smoker 2 | 52 | Female | 40 | 78 | ADC |
| Current Smoker 3 | 65 | Female | NA | 62 | LCC |
| Current Smoker 4 | 70 | Male | 36 | 72 | SCC |

**Supplementary Table 2. Bronchial tissue was obtained by lung resection from 4 never smokers and 4 current smokers at University Medical Center Groningen.** Lung resection was performed at University Medical Center Groningen (UMCG) on never and current smoker patients diagnosed with one of the following: lung adenocarcinoma (ADC), lung squamous cell carcinoma (SCC), large cell carcinoma (LCC), pulmonary carcinoid (PC), metastasis (MET), or inflammatory myofibroblastic tumour (IMT). Spirometry was also performed to assess lung function by measuring forced expiratory volume in one second (FEV1) relative to forced vital capacity (FVC).

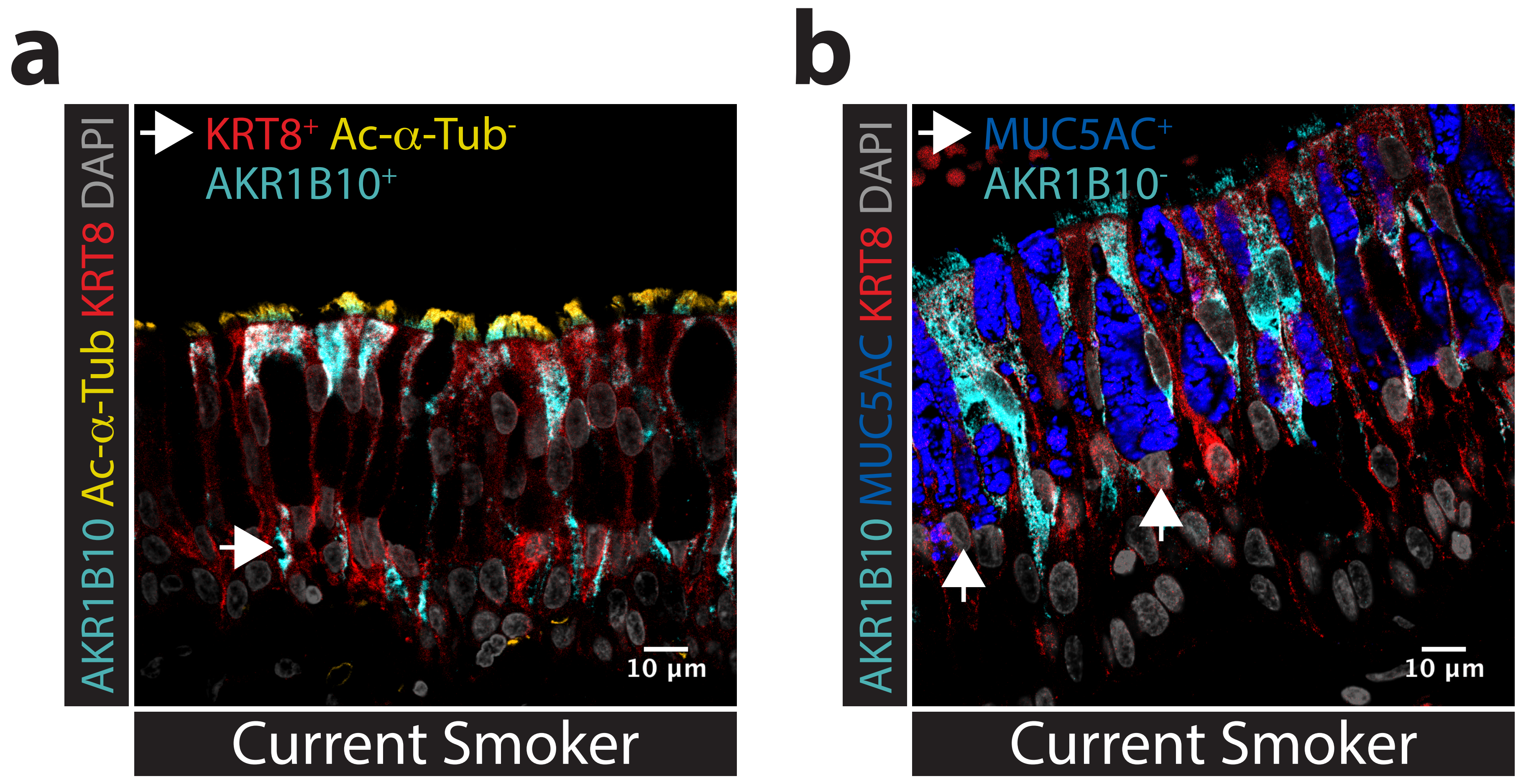


**Supplementary Figure 13. Non-ciliated cell AKR1B10 expression was uncommon.** (**a**) Immunostaining for AKR1B10, Acetylated alpha tubulin (Ac-α-Tub), and KRT8 was performed using bronchial tissue procured from an independent cohort of never and current smokers (UMCG cohort, **Supplementary Table 2**). A representative image of current smoker (right) tissue was displayed. An arrow was used to specify an example of a non-ciliated (Ac-α-Tub$^-$) KRT8$^+$ AKR1B10$^+$ cell. (**b**) Immunostaining for AKR1B10, MUC5AC (goblet), and KRT8 was also performed using bronchial tissue procured from an independent cohort of never and current smokers (UMCG cohort, **Supplementary Table 2**). A representative image of current smoker (right) tissue was displayed. Arrows were used to specify examples of goblet cells (MUC5AC$^+$) that did not express AKR1B10.

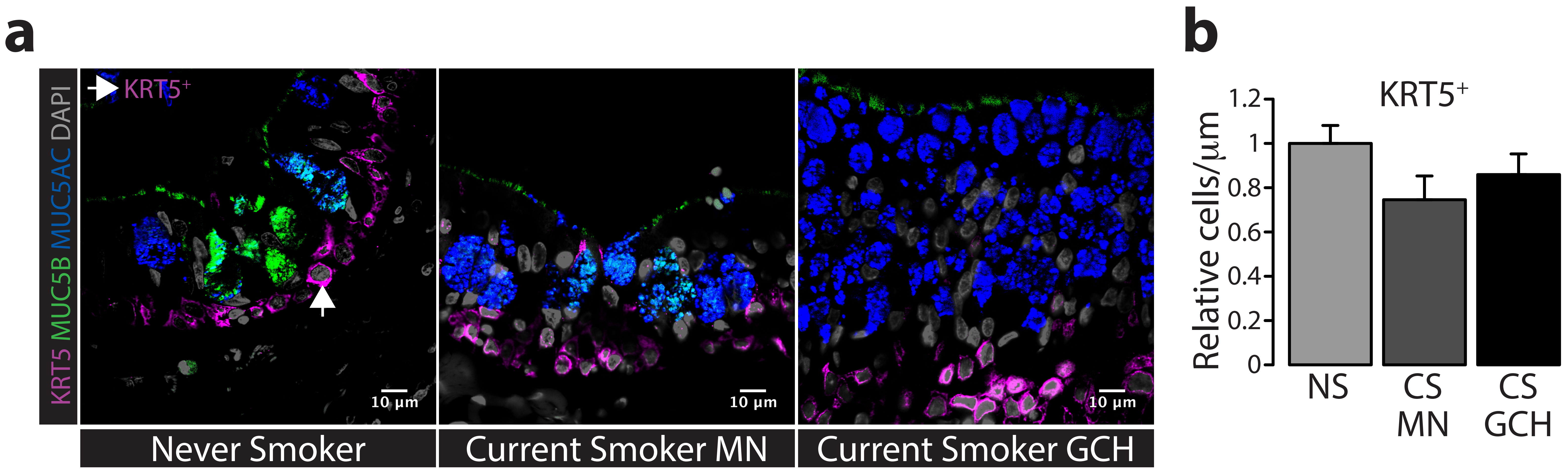


**Supplementary Figure 14. Basal cell cell numbers were not altered in smokers.** Immunostaining for KRT5, MUC5AC, and MUC5B was performed using bronchial tissue procured from an independent cohort of never and current smokers (UMCG cohort, **Supplementary Table 2**). (**a**) Representative images of never smoker tissue (left), as well as morphologically normal (MN) (middle) and goblet cell hyperplasia (GCH) (right) current smoker tissue were displayed. (**b**) Changes in tissue length (µm)-normalized numbers of KRT5$^+$ cells in MN and GCH current smoker tissue, relative to never smokers, were assessed by WRS test (MN $p = 0.2$; GCH $p = 0.2$).

| | Age | Sex | Pack years | Diagnosis |
|---|---|---|---|---|
| Never Smoker 1 | 33 | Female | 0 | PA |
| Never Smoker 2 | 63 | Male | 0 | ADC |
| Never Smoker 3 | 79 | Male | 0 | BC |
| Never Smoker 4 | 82 | Male | 0 | PC |
| Never Smoker 5 | 82 | Male | 0 | ADC |
| Current Smoker 1 | 76 | Female | 40 | ADC |
| Current Smoker 2 | 58 | Male | 40 | ADC |
| Current Smoker 3 | 61 | Male | 46 | SCC |
| Current Smoker 4 | 67 | Female | 45 | ADC |
| Current Smoker 5 | 71 | Male | 50 | ADC |

**Supplementary Table 3. Bronchial tissue was obtained by lung resection from 5 never smokers and 5 current smokers at University College London Hospital.** Lung resection was performed at University College London (UCL) Hospital on never and current smoker patients diagnosed with one of the following: lung adenocarcinoma (ADC), lung squamous cell carcinoma (SCC), basaloid carcinoma (BC), pulmonary carcinoid (PC), or pulmonary aspergilloma (PA). All specimens were procured at least 5 cm from bronchial sites affected by disease diagnoses and analyses indicated that tissue was histologically normal.

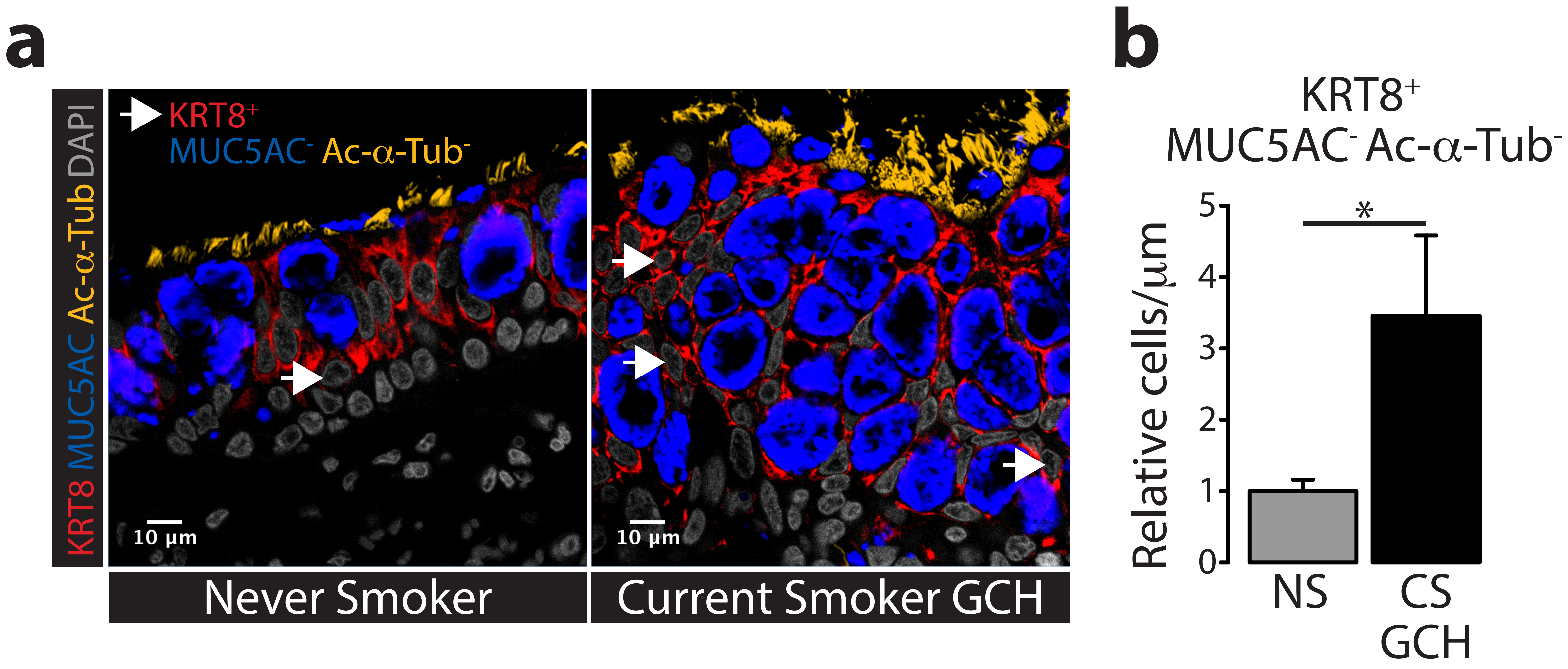


**Supplementary Figure 15. Increased numbers of indeterminate KRT8$^+$ cells were observed in goblet cell hyperplasia smoker tissue.** Immunostaining for KRT8, MUC5AC, and Acetylated alpha tubulin (Ac-α-Tub) was performed using bronchial tissue procured from an independent cohort of never and current smokers (UCL cohort, **Supplementary Table 3**). (**a**) Representative images of never smoker (left) and goblet cell hyperplasia (GCH) current smoker (right) tissue were displayed. Arrows were used to specify examples of KRT8$^+$ MUC5AC$^-$ Ac-α-Tub$^-$ cells. (**b**) Changes in tissue length (mm)-normalized numbers of KRT8$^+$ MUC5AC$^-$ Ac-α-Tub$^-$ cells in GCH current smoker tissue, relative to never smokers, were assessed by WRS test ($p = 0.04$).
